# Diversification of Dentate Gyrus Granule Cell Subtypes is Regulated by Neuregulin1 Nuclear Back Signaling

**DOI:** 10.1101/2024.10.15.618334

**Authors:** Prithviraj Rajebhosale, Li Jiang, Haylee J. Ressa, Kory R. Johnson, Niraj Desai, Alice Jone, Lorna W. Role, David A. Talmage

## Abstract

Neuronal heterogeneity is a defining feature of the developing mammalian brain, but the mechanisms regulating the diversification of closely related cell types remain elusive. In this study, we investigated the heterogeneity of dentate gyrus (DG) granule cells (GCs) and the influence of a psychosis associated V_321_L mutation in Neuregulin1 (Nrg1) on GC subtype composition. Using morpho-electric characterization, single-nucleus gene expression, and chromatin accessibility profiling, we identified distinct morphological and molecular features of typical GCs and a rare subtype known as semilunar granule cells (SGCs). The V_321_L mutation disrupts Nrg1 nuclear back-signaling, resulting in an overabundance of SGC-like cells. We discovered pseudotime gene expression trajectories suggesting the potential for GC-to-SGC transitions, supported by the accessibility of SGC-specific genes in other GCs. Intriguingly, we found an increase in SGC-marker expression over the adolescence to adulthood transition window in wild-type mice, coinciding with a decline in Nrg1 nuclear back-signaling capacity. This suggests that intact Nrg1 signaling suppresses SGC-like fate acquisition, and that its natural downregulation may underlie the emergence of SGC-like cells during postnatal development. Similarly, a pathological block of nuclear back signaling by the V_321_L mutation in Nrg1, may result in acquisition of the SGC-like fate due to loss of the repressive mechanisms maintained by intact nuclear back signaling. Our findings reveal a novel role for Nrg1 in maintaining DG cell-type composition and suggest that disrupted subtype regulation may contribute to disease-associated changes in DG GC morphology and function. Understanding these mechanisms provides new insights into mechanisms of cell-type diversity and its potential role in psychiatric pathology.

## Introduction

Single-cell genetic studies over the last decade have revealed that heterogeneity of neuronal cells is an emergent property of the developing mammalian brain (Tasic, Menon et al. 2016, Cembrowski and Spruston 2019, La Manno, Siletti et al. 2021, Network 2021, Yao, van Velthoven et al. 2023). However, how closely related neuronal cell types diversify remains mechanistically elusive (Morris 2019, Zeng 2022). In this study we comprehensively profiled differences between two closely related, yet morphologically distinct dentate gyrus (DG) granule cell (GC) types-typical GCs and semilunar GCs (SGCs), to investigate molecular determinants of individual cell type identity.

Generation of GCs in the DG proceeds in two main phases – embryonic and postnatal, which in mice, is further divided into a perinatal burst of neurogenesis (P0-P8), and “adult” neurogenesis (P8 into adulthood). Embryonically, GCs are produced from neural precursors (NPs) located in the dentate neuroepithelium (DNE) (Hatami, Conrad et al. 2018). These precursors migrate to establish a new neurogenic niche below the granule cell layer (GCL) known as the subgranular zone (SGZ) which gives rise to new GCs throughout life. Detailed transcriptomic analyses have concluded that the gene expression profiles of DNE and SGZ NPs do not significantly differ and that the neurogenic trajectories, i.e. sequential changes in the transcriptome as neurons differentiate, are also highly conserved between embryonic and postnatal development (Hochgerner, Zeisel et al. 2018, Berg, Su et al. 2019). However, a notable difference between embryonically and postnatally born GCs is that about half of the GCs produced during embryonic development are morphologically unique, bearing multiple splaying primary dendrites, known as semilunar granule cells (SGCs), which eventually comprise up to 3% of all DG GCs (Save, Baude et al. 2019). SGCs are not generated postnatally as evidenced by a spatial bias in their location within the GCL. SGCs are predominantly located in the suprapyramidal blade of the DG where they populate the dorsal most layers bordering the molecular layer (MOL) along with morphologically typical GCs which are born around the same time. Postnatally born GCs populate the GCL in an “outside-in” manner and are in cell layers closer to the hilus of the DG (Save, Baude et al. 2019). Studies characterizing the dendritic morphology of GCs and SGCs have concluded that the differences between these cells exists as early as 1-2 weeks after birth and persist thereafter, concluding that the SGC phenotype represents a bona fide cell type, supporting the conclusion that the two GC subtypes are fundamentally distinct cell types (Gupta, Proddutur et al. 2020).

The significance of understanding mechanisms related to heterogeneity of GCs is underscored by numerous findings of heterotopic and overabundant adult-born neurons with SGC-like morphologies in the DG of mice harboring disease-associated genetic mutations (Kim, Duan et al. 2009, Llorens-Martin, Jurado-Arjona et al. 2014). A similar phenotype has also been observed in postmortem brains of humans with schizophrenia, Alzheimer’s disease, and frontotemporal dementia(Lauer, Beckmann et al. 2003, Senitz and Beckmann 2003, Terreros-Roncal, Flor-Garcia et al. 2019, Marquez-Valadez, Rabano et al. 2022). These data imply potential conversion of GCs to SGCs or at the very least, a preserved capacity of the adult DG to generate SGC-like cells. Additionally, implicit in these findings is the idea that dysregulation of the cell biological mechanisms regulating subtype specification or maintenance might underlie the altered composition of the GC population in disease states.

In this study we performed morpho-electric and molecular characterization of GCs and SGCs to reveal cell-type specific features that distinguish GCs and SGCs, while also finding many similarities consistent with the close relatedness of these cells. We investigated this “relatedness” by performing trajectory analyses on gene expression and chromatin accessibility data obtained from GCs and SGCs and found the potential for a GC➔SGC genetic conversion implicated in aging and genetic associations with psychiatric disease. In line with this we found that a psychosis-associated missense mutation (Val_321_➔Leu; V_321_L) in Neuregulin1 (Nrg1) resulted in overabundant heterotopic SGCs, and that aging in the wild-type DG also resulted in overabundant and heterotopic cells with SGC-like gene expression. The V_321_L mutation diminishes the ability of Nrg1 proteins to perform membrane to nucleus signaling (nuclear back signaling), which was shown to regulate adult neurogenesis in the DG. We found that expression of requisite components of the Nrg1 nuclear back signaling pathway decreases over age. Thus, we conclude that intact Nrg1 nuclear back signaling suppresses the SGC-like fate, and loss of this signaling results in de-repression of the SGC-specific gene expression signature resulting in acquisition of SGC-like properties.

## Results

### Semilunar granule cells have unique morphological and electrical properties

Using Golgi staining we identified two morphologically distinct types of GCs distinguished by the number of primary dendrites (**Figure 1A**). Previously published reports have characterized a rare GC-like population known as semilunar granule cells (SGCs), which bear multiple splaying primary dendrites with cell bodies located near or within the inner molecular layer of the DG, and unique electrical properties (Cajal 1911, Williams, Larimer et al. 2007, Save, Baude et al. 2019, Gupta, Proddutur et al. 2020). Given the limitations of unambiguously resolving cell position within the granule cell layer (GCL) using Golgi staining, we performed whole cell patch clamp recordings from acute hippocampal slices with biocytin fills to reconstruct morphologies of recorded neurons, while simultaneously obtaining information regarding their intrinsic electrical properties (**Figures 1B & C**). We recorded GCs from the middle of the suprapyramidal blade of the DG and avoided the lower cell layer to avoid immature GCs generated from NSCs located in the SGZ. Similar to our results from Golgi staining, we found recorded and filled neurons with single and multiple primary dendrites that we classified as GCs and SGCs respectively (**Figures 1B-E**).

**Figure 1.**
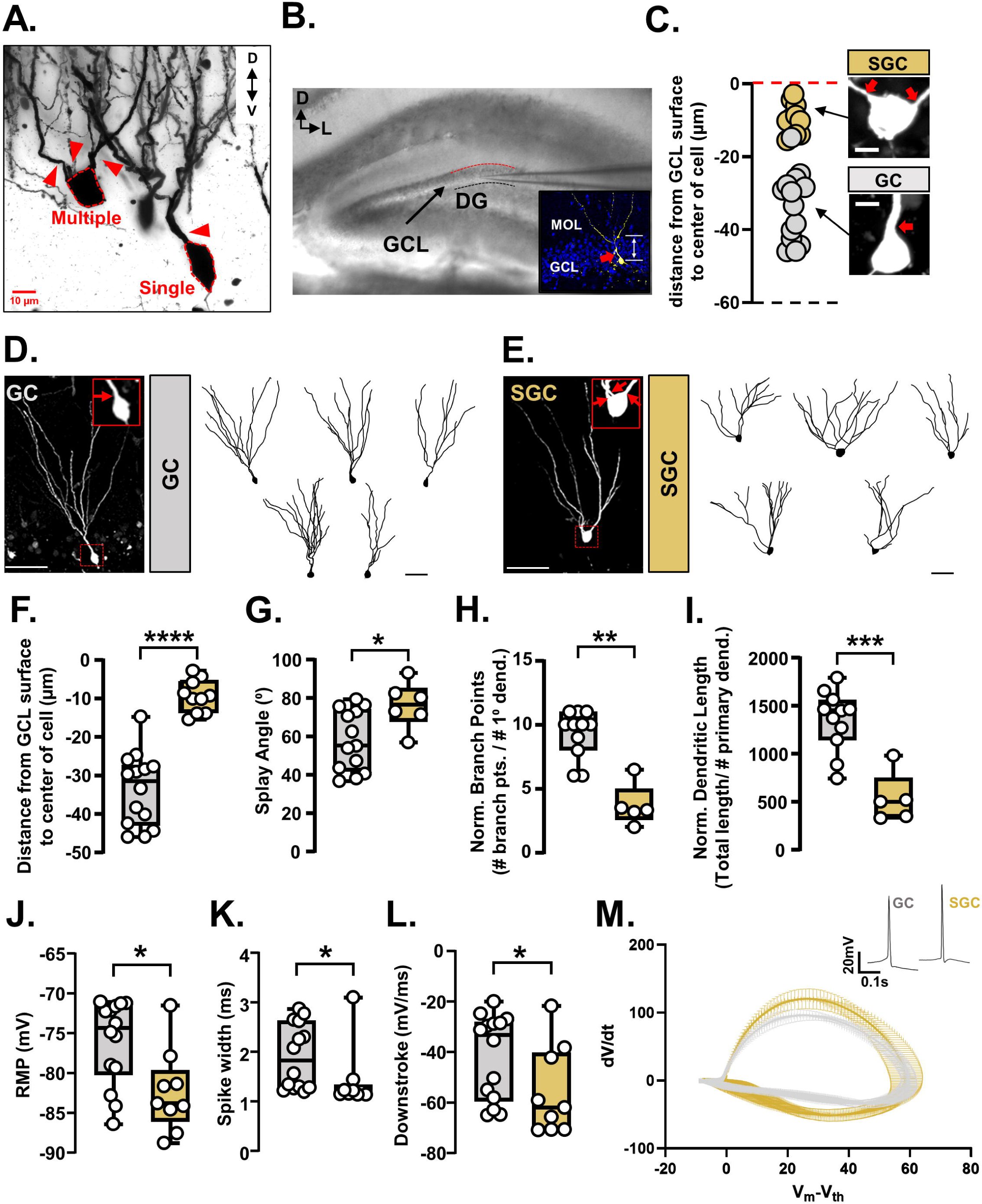
Morphological and electrical properties of semilunar granule cells are distinct from granule cells. **A.** Representative image of a WT Golgi-stained DG GCL showing two GCs, one example of a GC with a single primary dendrite and another of a GC with multiple primary dendrites. The cell body is outlined with a red dashed line. Primary dendrites are indicated by red arrowheads. Scale bar=10µm. **B.** DIC image of a coronal hippocampal section prepared for acute slice electrophysiology. The anatomical legend in the top left corner indicates the orientation of the slice – D= dorsal and L=lateral. The dorsal edge of the granule cell layer (GCL) is demarcated by a red dashed line, and the ventral border by a black dashed line. The internal solution within the patch pipette contained biocytin to fill the recorded cells. Scale bar=100µm. DG= dentate gyrus, GCL= granule cell layer. (Inset) A representative confocal image of a patched granule cell filled with biocytin identified via fluorescently conjugated streptavidin. The image shows a maximum z-projection of three 1µm optical sections containing the widest portion of the soma used to quantify cell position in the GCL relative to the GCL-MOL boundary (red dashed line). Scale bar= 50µm. MOL= molecular layer. **C.** Quantified cell positions of all recorded GCs from WT mice. The red and black dashed lines correspond to the dorsal and ventral borders of the GCL respectively. Semilunar granule cells (SGCs - GCs with multiple primary dendrites) were found in the top cell body layer of the GCL (gold circles). Typical GCs with a single primary dendrite were found throughout the GCL (silver circles). Representative images of a filled and recorded SGC and GC are shown. Red arrows point to primary dendrites extending from the cell body. N= 14 cells (GC) and 10 cells (SGC) from 5 mice (20 slices). Scale bar= 10µm. **D.** (**Left**) Representative image of a filled GC. Dotted red box indicates the region magnified in the inset. Inset shows a magnification of the cell body and proximal dendrite of the filled GCs. Red arrow marks the primary dendrite emanating from the soma. Scale bar= 50µm. (**Right**) Five representative 2-D tracings of filled GCs. Scale bar= 50µm. **E.** (**Left**) Representative image of a filled SGC. Dotted red box indicates the region magnified in the inset. Inset shows a magnification of the cell body and proximal dendrites of the filled SGCs. Red arrows mark primary dendrites emanating from the soma. Scale bar= 50µm. (**Right**) Five representative 2-D tracings of filled GCs. Scale bar= 50µm. **F.** Quantification of cell body position within the GCL of GCs (silver) and SGCs (gold). SGC cell body locations were significantly different from those of GCs (****p<0.0001, Welch’s t-test (two tailed); t=8.471, df=19.16). N=14 cells (GC) and 10 cells (SGC) from 5 mice. **G.** Quantification of splay angle of GCs and SGCs. The splay angle (angle between the two farthest dendrites measured at the center of the cell body) was significantly larger in SGCs compared to GCs (*p=0.0168, Welch’s t-test (two tailed); t=2.754, df=12.59). N=13 cells (GC) and 6 cells (SGC) from 4 mice. **H.** Quantification of dendritic branching of GCs and SGCs normalized to the number of primary dendrites. The number of branch points was significantly higher in GCs compared to SGCs (**p=0.0023, Mann-Whitney, U=2). N=11 cells (GC) and 5 cells (SGC) from 4 mice. **I.** Quantification of dendritic length of GCs and SGCs normalized to the number of primary dendrites. The normalized dendritic length was significantly higher in GCs compared to SGCs (***p=0.0004, Welch’s t-test (two tailed); t=5.348, df=9.366). N=11 cells (GC) and 5 cells (SGC) from 4 mice. **J.** SGCs have a significantly more hyperpolarized resting membrane potential compared to GCs (*p=0.02, Mann-Whitney, U=26; N= 14 cells (GCs) and 9 cells (SGCs) from 5 mice). **K.** SGCs have a significantly narrower spike width compared to GCs (*p=0.012, Mann-Whitney, U=24; N= 14 cells (GCs) and 9 cells (SGCs) from 5 mice). **L.** SGCs have a significantly faster downstroke compared to GCs (*p=0.046, Mann-Whitney, U=31; N= 14 cells (GCs) and 9 cells (SGCs) from 5 mice). **M.** Phase plots of action potential dynamics (dV/dt vs. V_m_-V_th_) provide a visualization of differences in the AP waveform between GCs and SGCs. N= 14 cells (GCs) and 9 cells (SGCs) from 5 mice. <u>Note</u>: the difference in number of GCs and SGCs for different morphological and electrical measures was due to incomplete total reconstructions which precluded inclusion for dendritic morphology parameters.

In the DG of WT mice, we found that SGCs were in the top cell body layers (<20µm from the surface) and cells with the typical GC morphology were found throughout the GCL (**Figures 1F**, Welch’s t-test (two-tailed) p<0.0001, t=8.471, df=19.16). SGCs had wider dendritic splay angles (**Figure 1G**, Welch’s t-test (two-tailed) p=0.02, t=2.754, df=12.59) and less complex dendrites (**Figure 1H**, branch points per dendrite, Mann-Whitney test p=0.002 U=2; **Figure 1I**, length per dendrite, Welch’s t-test (two-tailed) p=0.0004, t=5.348, df=9.366). These findings match existing reports regarding morphologies of embryonically born GCs (Kerloch, Clavreul et al. 2019).

We recorded the membrane voltage responses of GCs and SGCs to current steps. Action potentials (APs) at rheobase were analyzed for twenty features. We found that SGCs had a significantly hyperpolarized resting membrane potential (RMP) (**Figure 1J**, Mann-Whitney Test p=0.02 U=26), shorter AP width (**Figure 1K**, Mann-Whitney Test p=0.01 U=24), and faster downstroke (**Figure 1L**, Mann-Whitney Test p=0.046 U=31). The pronounced differences in the kinetics of the SGC action potentials can be seen in the phase plots for APs at rheobase (**Figure 1M**).

Since SGCs are born earlier than most other GCs, we asked if the group differences between GCs and SGCs could be reflective of cellular age as opposed to cell-type defining features. Given the stereotyped relationship between GCL lamination and neuronal birthdate in the DG, we analyzed the relationships between the measured electrical properties and soma position within the GCL. We found that RMP (**Figure S1A**, p=0.0129), spike width (**Figure S1B**, p=0.0543), and downstroke (**Figure S1C**, p-0.0139) significantly co-varied with soma position as expected from the GC vs. SGC grouped analyses (**Figure 1J-L**). We also found that rheobase (**Figure S1D**, p=0.0475), sag (**Figure S1E**, p=0.0359), and upstroke-downstroke ratio (**Figure S1F**, p=0.0131) significantly co-varied with soma positions in the GCL. The remainder of the properties did not differ significantly between GCs and SGCs or by soma position.

Next, we assessed the correlations between these six properties and soma positions <u>for GCs alone</u> (excluding SGCs) to identify electrical properties that might reflect effects of cell birthdate rather than cell type. Of the six properties, we found that downstroke and sag were still significantly correlated with soma positions (**Figure S1I** downstroke p=0.006, and **Figure S1K** sag p=0.03), while the rest of the properties did not correlate with soma positions (**Figure S1G** RMP p=0.22, **Figure S1H** spike width p=0.1, **Figure S1J** rheobase p=0.06, and **Figure S1L** upstroke-downstroke ratio p=0.4). These results indicate that the differences between GCs and SGCs in RMP and spike width are cell type defining features.

Thus, our findings define electrophysiological differences between GCs and SGCs further substantiating published reports of electrophysiological differences between GCs and SGCs (Williams, Larimer et al. 2007, Kerloch, Clavreul et al. 2019).

### V_321_L mutation in Nrg1 results in altered composition of the DG granule cell population

Cells with SGC-like morphologies are overabundant in postmortem brains of people with neuropsychiatric conditions and in rodent experimental systems modeling psychiatric disease-associated genetic and environmental factors (Lauer, Beckmann et al. 2003, Senitz and Beckmann 2003, Kim, Duan et al. 2009, Fitzsimons, van Hooijdonk et al. 2013, Llorens-Martin, Jurado-Arjona et al. 2014, Howe, Li et al. 2017, Terreros-Roncal, Flor-Garcia et al. 2019, Marquez-Valadez, Rabano et al. 2022). We previously characterized a psychosis-associated missense mutation in the *NRG1* gene showing that it resulted in reduced stem cell proliferation, increased neuronal differentiation, and altered dendritic branching of GCs in the DG (Rajebhosale, Jone et al. 2024). During this study, we qualitatively noted numerous GCs with multiple primary dendrites resembling SGCs in the V_321_L DG. To quantify the numbers of GCs and SGCs, we used the rapid Golgi staining technique and counted stained neurons with single or multiple primary dendrites from WT and V_321_L DG (**Figure 2A**). There was no difference in the total number of cells labeled by the Golgi stain between genotypes (**Figure 2A, left;** Welch’s t-test (two-tailed) p=0.68, t=0.4338, df=6.972). V_321_L mice had a significantly higher number of GCs with multiple primary dendrites compared to WT mice and a significantly lower number of GCs with single primary dendrites compared to WT mice **(Figure 2A, right;** Two-way ANOVA Genotype x #Primary dendrites p=0.011, F (1,7) = 11.81. Multiple comparisons (Bonferroni corrected): WT multiple vs. V_321_L multiple p=0.008; WT single vs. V_321_L single p=0.008). The number of GCs with multiple primary dendrites also significantly differed from the numbers of GCs with single primary dendrites in the V_321_L DG (V_321_L single vs. V_321_L multiple, p=0.013), whereas there was no difference in this regard in WT mice (p=0.85). These results indicate that V_321_L mice have a higher number of cells with SGC-like morphologies in the DG, potentially at the expense of the other subpopulation of GCs.

**Figure 2.**
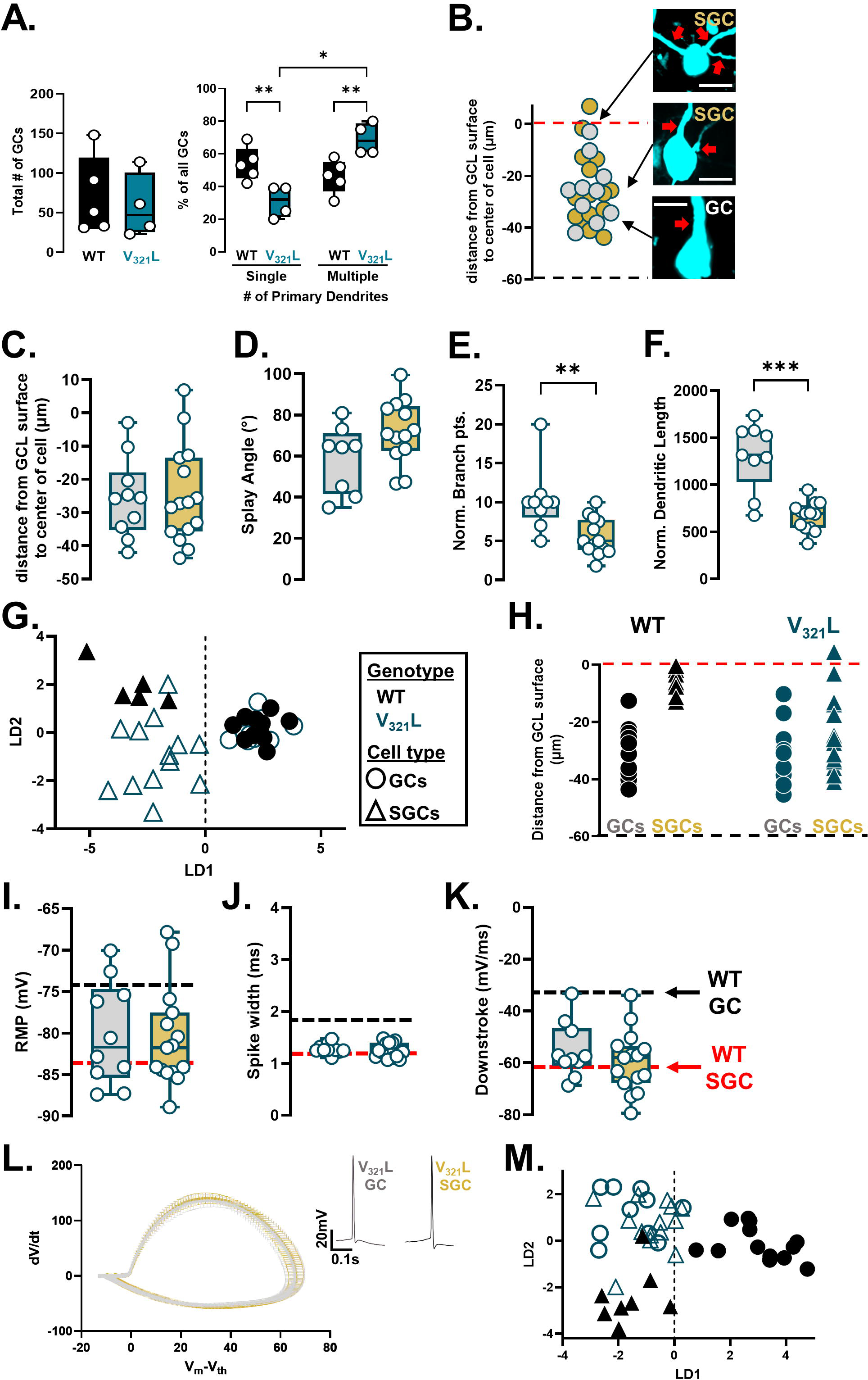
GCs in the V_321_L DG show SGC-like electrical properties. **A.** (**Left**) Quantification of total number of GCs from Golgi-stained DG, quantified from WT (black) and V_321_L (teal) DG. N=23-148 neurons/mouse from 5 mice. Each dot is total number of neurons from 1 mouse. (Welch’s t-test (two-tailed) p=0.7 t=0.4338, df=6.972) (**Right**) Quantification of percentage of GCs with single or multiple primary dendrites in WT and V_321_L DG. N=23-148 neurons/mouse from 5 mice (Two-way ANOVA Genotype x #Primary dendrites p=0.011 F (1,7) = 11.81; V_321_L single vs. V_321_L multiple *p=0.013. WT multiple vs. V_321_L multiple **p=0.008. WT single vs. V_321_L single **p=0.008). <u>Note</u>: The Golgi staining method is “non-random” and biases for labeling of superficially located GCs resulting in disproportionate labeling of SGCs. **B.** Quantified cell positions of all recorded GCs from V_321_L mice. The red and black dashed lines correspond to the dorsal and ventral borders of the GCL respectively. Both SGCs (gold circles) and GCs (silver circles) were found throughout the GCL. Representative images of filled and recorded SGCs and GC are shown. Red arrows point to primary dendrites extending from the cell body. N= 10 cells (GC) and 15 cells (SGC) from 5 mice (17 slices). Scale bar= 10µm. **C.** Quantification of cell body position within the GCL of GCs (silver) and SGCs (gold). SGC cell body locations are not significantly different to those of GCs (p=0.9012, Welch’s t-test (two tailed); t=0.1256, df=21.78; N=10 cells (GC) and 15 cells (SGC) from 5 mice). **D.** Quantification of splay angle of GCs and SGCs. The splay angle was not significantly different between GCs and SGCs (p=0.0811, Welch’s t-test (two tailed); t=1.879, df=14.11; N=8 cells (GC) and 13 cells (SGC) from 5 mice). **E.** The number of normalized branch points was significantly higher in GCs compared to SGCs (**p=0.0013, Mann-Whitney, U=13; N=9 cells (GC) and 13 cells (SGC) from 5 mice). **F.** The normalized dendritic length was significantly higher in GCs compared to SGCs (***p=0.0005, Welch’s t-test (two tailed); t=5.023, df=10.05; N=9 cells (GC) and 13 cells (SGC) from 5 mice). **G.** Linear discriminant analysis (LDA) of morphological properties of WT (black) and V_321_L (teal) cells reveals separation of GCs (circles) and SGCs (triangles) along the LD1 axis. **H.** V_321_L SGCs are localized throughout the GCL unlike WT SGCs. Each symbol represents a single cell from biocytin-filled reconstructions replotted from Figures 2B and 1C - WT (black) and V321L (teal); GCs (circles) and SGCs (triangles). Red dashed line represents the dorsal edge of the GCL. **I.-K.** V_321_L GCs and SGCs did not show significant differences in their electrophysiological properties. **I.** RMP (p=0.8374, Welch’s t-test. t=0.2081, df=18.86). **J.** Spike width (p=0.2776, Mann-Whitney U=55). **K.** Downstroke (p=0.5392, Mann-Whitney U=63.5). Black dashed line indicates the median values for <u>WT</u> GCs and red dashed line indicates median values for <u>WT</u> SGCs. N= 10 cells (GC) and 15 cells (SGC) from 5 mice. **L.** Phase plots of action potential dynamics (dV/dt vs. Vm-Vth) provide a visualization of the AP waveform of GCs and SGCs. N= 10 cells (GCs) and 15 cells (SGCs) from 5 mice. Inset shows representative action potential waveforms of a GC and SGC recorded at rheobase. **M.** Linear discriminant analysis (LDA) of electrical properties of WT (black) and V321L (teal) cells reveals separation of WT GCs (circles) from all other cells from both genotypes along the LD1 axis.

We next profiled GCs from V_321_L mice throughout the GCL for their electrical and morphological properties using whole cell patch clamp with dye fills. Like the WT DG, we found SGCs in superficial layers and GCs throughout the GCL. However, we also found cells with multiple primary dendrites throughout the GCL in the V_321_L DG (**Figure 2B**). Analyzing the locations of all recorded cells, we found no difference between GC and SGC localization (**Figure 2C**, Welch’s t-test (two-tailed) p=0.9, t=0.1256, df=21.78). Additionally, GCs from V_321_L mice showed wider splay angles than WT GCs, and there were no significant differences between GCs and SGCs (**Figure 2D**, Welch’s t-test (two-tailed) p=0.08, t=1.879, df=14.11). In contrast, the differences in dendritic complexity between GCs and SGCs were preserved in the V_321_L mutant DG (**Figure 2E**, normalized branch points, Mann-Whitney Test p=0.0013, U=13; **Figure 2F**, normalized dendritic length, Welch’s t-test (two-tailed) p=0.0005, t=5.023, df=10.05). To further assess any morphological changes in the V_321_L DG, we subjected the 15 measured morphological parameters from WT and V_321_L cells to classification using linear discriminant analysis (LDA). LDA resulted in successful capture of cell type differences between GCs and SGCs (along LD1) but not between genotypes (**Figure 2G**). Analysis of the weighted contributions of each morphological parameter to the LDs revealed that separation along LD1 was heavily influence by dendritic complexity parameters in line with the known lower complexity of SGC dendrites (**Figure S2A**). We also performed unsupervised clustering using k-means clustering which also resulted in clustering of cells primarily based on cell type rather than genotype (**Figure S2B**). These data indicate that although the SGC-like cells were over-represented and mis-localized in the V_321_L, they were morphologically indistinguishable from WT SGCs (**Figure 2H**).

We profiled sub- and suprathreshold intrinsic electrical properties of cells in the V_321_L DG. Unlike WT GCs and SGCs, we found no differences in electrical properties between V_321_L GCs and SGCs (**Figure 2I-K**, RMP: Mann-Whitney Test p=0.9, U=73; spike width: Mann-Whitney Test p=0.3, U=55; downstroke: Mann-Whitney Test p=0.5, U=63.5). The phase plots for APs at rheobase further demonstrate this point showing a virtually complete overlap between GCs and SGCs (**Figure 2L**).

To further evaluate effects of cell type and genotype we performed LDA and unbiased clustering using k-means clustering using all recorded features from WT and V_321_L neurons. LDA resulted in separation of WT GCs and all other groups (WT SGCs, V_321_L GCs and V_321_L SGCs) along LD1 driven strongly by spike width (**Figure 2M and Figure S2C**). k-means clustering also showed that V_321_L GCs and SGCs showed a nearly similar distribution as WT SGCs (**Figure S2D**). These data indicate that in the V_321_L mutant DG, SGCs are morpho-electrically “normal” and that morphologically typical GCs assume electrical properties of SGCs.

Thus, V_321_L mice have more cells with SGC-like morphologies and electrical properties. These data indicate a potential overabundance of SGCs in the V_321_L DG.

### V_321_L mice have a higher number and altered positioning of Penk+ GCs compared to WT mice

Recent studies have shown that GCs expressing the gene proenkephalin (*Penk*), have multiple primary dendrites and localization consistent with those of embryonically born GCs (Erwin, Sun et al. 2020, Mortessagne, Cartier et al. 2024). Using fluorescent in situ hybridization (FISH), we detected *Penk* transcripts in the DG of young adult mice and quantified the distance of *Penk*+ cells in the suprapyramidal granule cell layer (GCL) from the GCL-IML boundary (**Figure 3A**). In WT mice, the majority of the *Penk*+ GCs were in the top 2 cell body layers of the suprapyramidal GCL whereas this bias towards the GCL-IML boundary was lost in the V_321_L DG (**Figure 3B**; K-S Test p=0.013 D=0.36).

**Figure 3.**
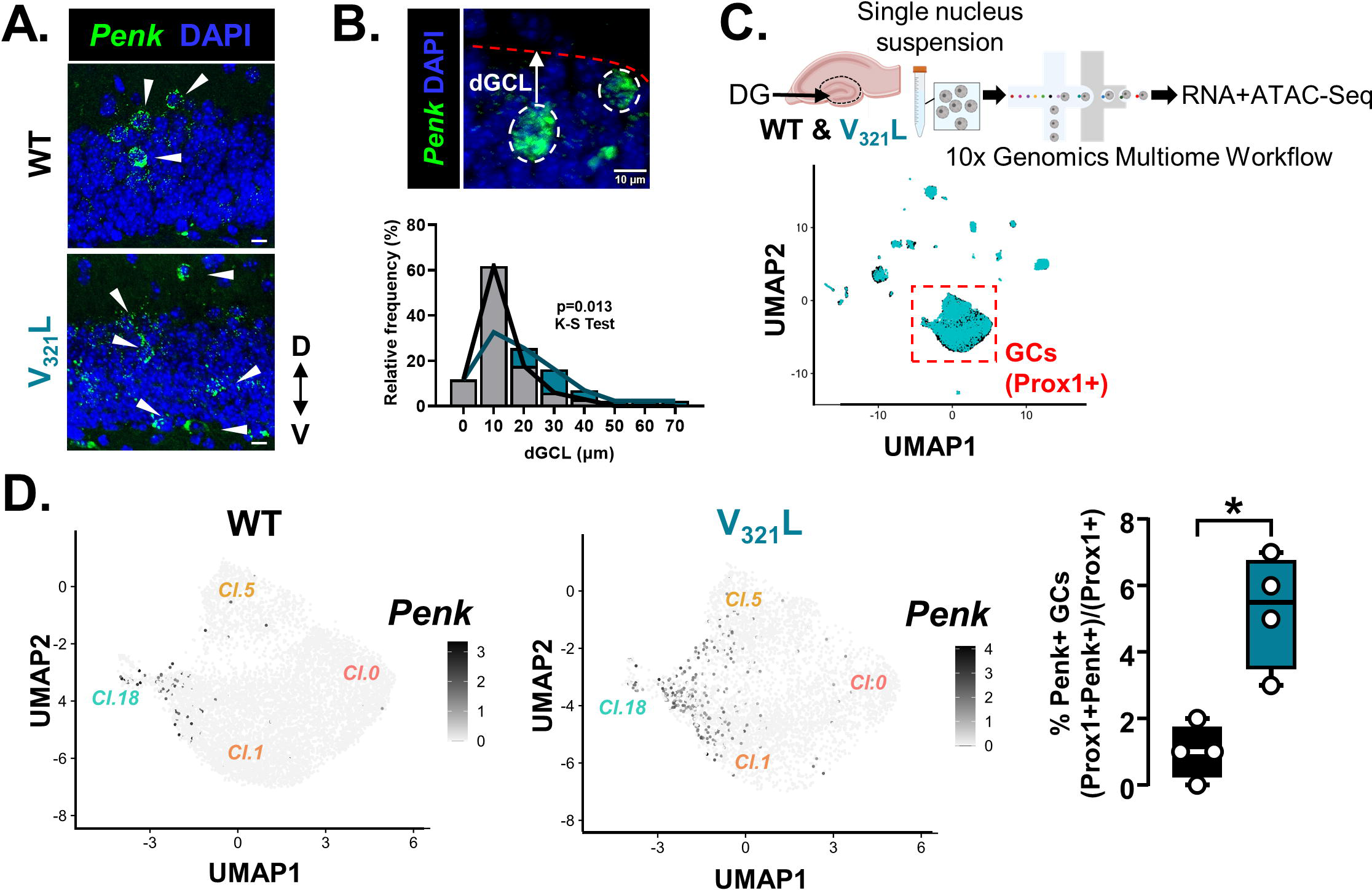
V_321_L mice have a higher number and altered positioning of Penk+ GCs compared to WT mice. **A.** (**Top**) Representative image of *Penk* mRNA (green) expression in the DG of a WT mouse. (**Bottom**) Representative image of *Penk* mRNA (green) expression in the DG of a V_321_L mouse. White arrowheads indicate Penk+ cells in the GCL and IML. Anatomical guide indicates dorsal (D) and ventral (V) directions. **B.** (**Top**) Representative image showing two *Penk*+ GCs outlined with white dashed lines. White arrow denotes distance of the *Penk*+ cell from the GCL-IML boundary (dGCL). Red dotted line delineates the GCL-IML boundary of the suprapyramidal blade (SPB) of the DG. Scale bar=10µm. (**Bottom**) Quantification of the soma positions within the GCL of the *Penk*+ cells from WT (black) and V_321_L (teal) mice. The distribution of *Penk*+ cells in the V_321_L DG was significantly altered compared to WT DG (Kolmogorov-Smirnov Test p=0.013, D=0.36). N=3 mice/genotype. **C.** (**Top**) Schematic of the workflow for single nucleus RNA- and ATAC-Seq (Multiome) of WT and V_321_L DG. (**Bottom**) UMAP of snRNA-Seq showing clusters of WT (black) and V_321_L (teal) nuclei. Red dashed box denotes clusters of GCs identified as *Prox1*+ excitatory neurons (**See Figure S3A and S3B**). N=4 mice/genotype; 2 male and 2 female. **D.** UMAPs showing expression of *Penk* in GCs of WT (**Left**) and V_321_L mice (**Middle**). The level of expression is indicated by the heatmap alongside. (**Right**) Quantification of percentage of GCs expressing *Penk*+ in WT and V_321_L mice showing that V_321_L mice had significantly more *Penk*+ GCs (Mann-Whitney Test (two-tailed) *p=0.03, U=0). N=4 mice/genotype.

To examine molecular distinctions between GC subtypes, we performed single nucleus RNA and ATAC sequencing of single nucleus suspensions prepared from WT and V_321_L microdissected DGs (**Figure 3C, top**). We detected all major cell types across both genotypes indicating no overt differences in clustering or cell composition of the DG between WT and V_321_L mice (**Figure 3C, bottom; Figure S3A; Figure S4A-O**). We found no differences in the total number of *Prox1*+ neurons (GCs) (**Figure S3B;** Mann-Whitney Test p=0.69, U=6). Upon examination of *Penk* expression in the GC clusters, we found that *Penk*-expressing cells subclustered within the larger clusters of GCs, and that the total percentage of *Penk*+ GCs was significantly higher in V_321_L mice (**Figure 3D**; Mann-Whitney Test p=0.03, U=0). We found 4 clusters of GCs – cluster 0,1,5, and 18. While a greater absolute number of *Penk*+ GCs were found in cluster 1 in proximity to cluster 18, cluster 18 contained the highest percentage of *Penk*+ cells indicating enrichment for SGCs in cluster 18 in both genotypes (**Figure S3C**). Marker gene discovery for the different GC clusters revealed relatively minor distinctions in gene expression between clusters 0,1 and 5. However, cluster 18 was uniquely marked by expression of Sortilin Related VPS10 Domain Containing Receptor 3 (*Sorcs3*) (**Figure S3D, left**). We performed FISH for *Sorcs3* transcripts in the DG and found a spatial bias of *Sorcs3*+ cells towards the GCL-IML boundary like *Penk*+ cells (and SGCs) consistent with their embryonic origins (**Figure S3D, right**).

### Discovery of novel marker genes for semilunar granule cells

We observed greater numbers and altered localization of *Penk*+ GCs in the V_321_L DG (**Figures 3A, B & D**). We identified *Sorcs3* as a marker for GCs located in the Penk-enriched GC cluster (cl.18) (**Figure S3D**). Thus, we next asked if V_321_L mice had greater numbers and/or altered localization of *Penk and Sorcs3* co-expressing GCs using FISH. We found that V_321_L mice had significantly higher numbers of cells co-expressing *Penk* and *Sorcs3* (**Figure 4A**, Welch’s t-test (two-tailed) p=0.014, t=5.278, df=2.941). We also found a significantly higher number of Penk and Sorcs3 co-expressing cells in V321L mice using snRNASeq (**Figure 4b’’**, Mann-Whitney Test p=0.03, Two-stage step-up Benjamini-Krieger-Yekutieli FDR corrected for multiple comparisons q=0.002, U=0). Additionally, we noted that the *Penk+Sorcs3+* cells in the V_321_L DG were not restricted to the GCL-IML boundary cell layers (**Figure 4a’**, Kolmogorov-Smirnov test p=0.029, D=0.4), much like the morphologically identified SGCs (**Figure 2H**).

**Figure 4.**
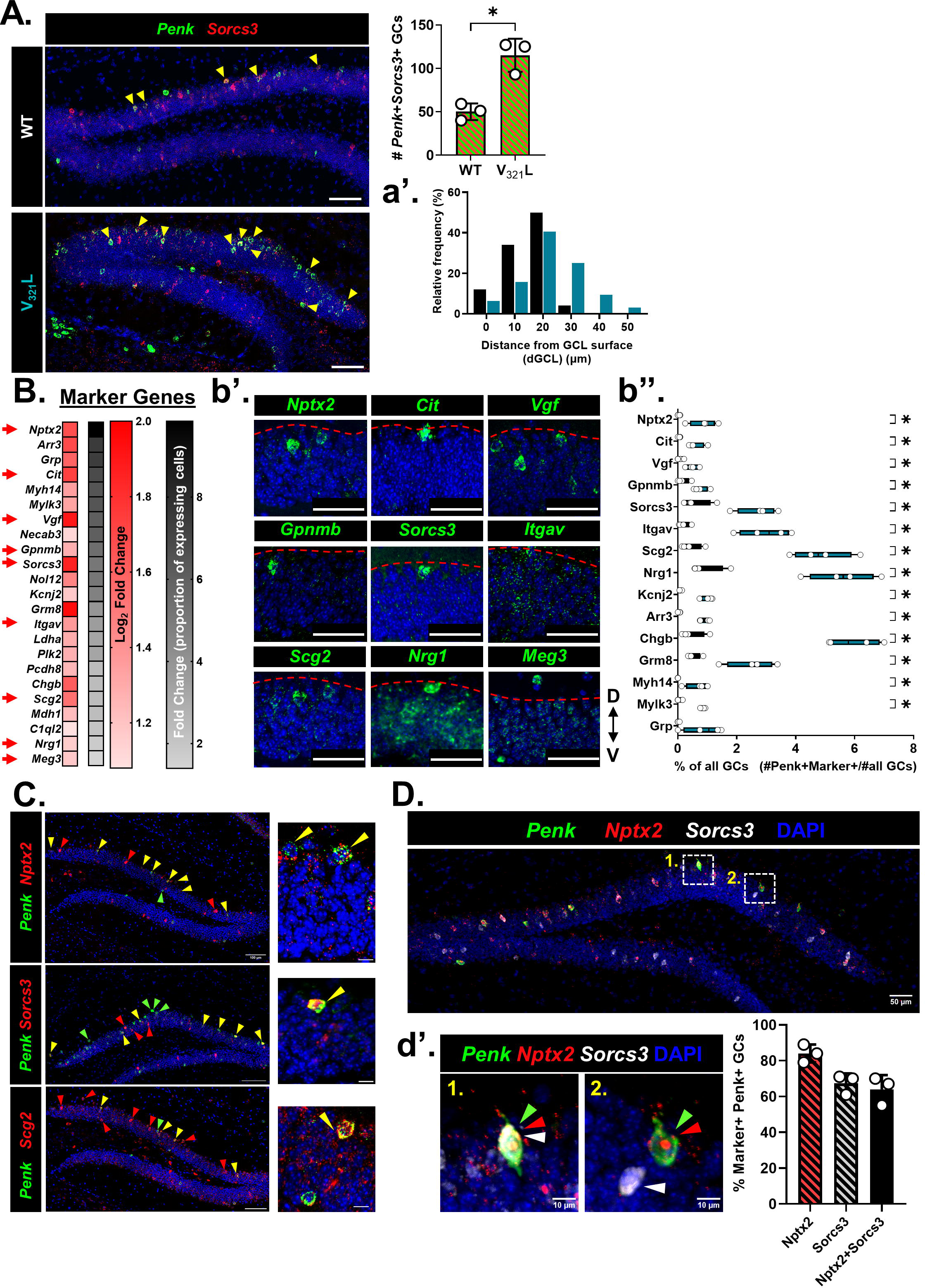
Discovery of novel marker genes for semilunar granule cells. **A.** Co-expression of *Penk* and *Sorcs3* was evaluated in the V_321_L DG. Representative image from a 1-month-old WT and V_321_L DG showing cells expressing *Penk* (green), *Sorcs3* (red). Yellow arrowheads indicate *Penk+Sorcs3+* cells. Scale bar= 100µm. (**Right**) Quantification of number of *Penk+Sorcs3+* cells in the suprapyramidal blade (SPB) of the DG of WT and V_321_L mice. N= 9 sections from 3 mice, each dot represents one mouse. There were significantly more *Penk+Sorcs3+* cells in the V_321_L DG (Welch’s t-test (two-tailed) p=0.014, t=5.278, df=2.941). **a’.** Frequency distribution of location (dGCL) of *Penk+Sorcs3+* cells in the SPB of the DG of WT and V_321_L mice (N=3 mice each). The spatial distribution of labeled cells was significantly different between the genotypes, with the V_321_L mice showing more Penk+Sorcs3+ cells present in the lower layers of the GCL, away from the GCL-IML boundary (K-S test, p=0.029, D=0.4). **B.** Marker gene discovery for *Prox1*+*Penk*+ GCs (SGCs) identified several genes whose expression was enriched in SGCs. The red gradient heatmap shows expression of the designated genes as Log2 fold change relative to non-SGCs (*Prox1*+*Penk*-GCs). The accompanying black gradient heatmap indicates the degree of exclusivity of expression of the identified putative SGC marker genes (% *Penk*+ GCs/ % *Penk*-GCs). Red arrows indicate genes chosen for validation via ISH. **b’.** Expression of the indicated subset of the putative marker genes was evaluated in the young adult DG (2 months) using RNAScope ISH. Images of the middle of the GCL in the suprapyramidal blade of the DG were examined. Most of the genes enriched in *Penk*+ GCs were found to be highly expressed in cells located closer to/ at or beyond the boundary between the GCL and IML (red dashed line) in line with stereotypical location of SGCs. Scale bar= 50µm. **b’’.** Quantification of % of *Penk*+ GCs expressing selected marker genes in WT and V_321_L mice. Number of cells co-expressing *Penk* and each of the marker genes were obtained from the snRNASeq data and divided by the total number of cells in clusters 0,1,5,18. WT and V_321_L samples were compared for each gene using multiple Mann-Whitney tests with an FDR correction. With the exception of Grp (WT vs. V321L FDR adj.p=0.14, U=2.5), V321L samples had significantly more cells co-expressing Penk and selected marker genes (WT vs. V321L FDR adj.p=0.03, U=0) (N=4 mice/genotype; 2M, 2F). **C.** Co-expression of *Penk* (green) and several marker genes (red) was evaluated using RNAScope multiplex ISH. Images showing co-expression of Penk with either Nptx2 (**top**), Sorcs3 (**middle**), or Scg2 (**bottom**) are shown. Scale bar= 100µm. Red arrowheads indicate cells expressing marker gene alone, green arrows indicate cells expressing *Penk* alone, and yellow arrows indicate cells co-expressing *Penk* and marker gene. Inset shows a higher magnification image of a few example cells with *Penk*-marker co-expression (yellow arrowheads). Scale bar= 10µm. **D.** Co-expression of *Penk*, *Nptx2*, and *Sorcs3* was evaluated as a defining feature for identification of SGCs. Representative image from a 2-month-old WT DG showing cells expressing *Penk* (green), *Nptx2* (red), and *Sorcs3* (white). Scale bar= 50µm. Insets labeled 1 and 2 show cells expressing different combinations of the selected markers. Inset 1 shows a *Penk+Nptx2+Sorcs3*+ cell. **d’.** (**Left**) Inset 2 shows two labeled cells-one expressing *Sorcs3* alone (white arrowhead), and another expressing *Penk* and *Nptx2* (green and red arrowheads). Scale bar= 10µm. (**Right**) Quantification of percentage of *Penk*+ cells expressing the selected marker gene combinations in the suprapyramidal blade of the DG of WT mice. N= 9 sections from 3 mice, each dot represents one mouse. Percentage of *Penk*+*marker*+ cells is as follows (mean ± std.dev): *Nptx2* (red with black patterned fill) 84±5%, *Sorcs3* (grey with black patterned fill) 67.3±5.5%, and *Nptx2+Sorcs3* (black filled bar) 64±7.9%.

To ascertain whether the increase in *Penk*+ or *Penk+Sorcs3+* GCs reflected upregulation of *Penk* and/or *Sorcs3* due to loss of Nrg1 nuclear back signaling, or whether it indeed reflected an increase in molecularly defined SGCs, we sought to identify additional genetic markers for this cell type. We reasoned that combinatorial detection of such markers could discern changes in cell type composition vs. changes in regulation of *Penk* expression. We assessed differential gene expression in *Penk*+ GCs compared to *Penk*-negative GCs yielding 102 genes whose expression was significantly enriched in *Penk*+ GCs (fold change of ≥2 and adj. p≤0.05). We rank ordered these genes based on the ratio of percentage of *Penk*+ cells expressing them to the percentage of *Penk*-cells expressing them; 23 of these genes are displayed in **Figure 4B** (See **Table S1**). Of these, we selected 9 genes for further validation using FISH to assess spatial localization consistent with embryonically born GCs. We found that different genes displayed varying degree of spatial bias towards the GCL-IML boundary with some showing a more exclusive spatial segregation such as Neuronal pentraxin 2 (*Nptx2),* Citron (*Cit*), and *Sorcs3*, which we showed earlier to also be a marker gene for cluster 18-the cluster enriched for *Penk*+ GCs (**Figure 4b’**). We next quantified the percentage of *Penk*+ GCs co-expressing each of these marker genes in the WT and V_321_L DG snRNASeq datasets. Except for gastrin releasing peptide (*Grp*) (Mann-Whitney Test p=0.1, Two-stage step-up Benjamini-Krieger-Yekutieli FDR corrected for multiple comparisons q=0.01, U=2.5), all other marker gene-Penk co-expressing cells formed a significantly higher proportion of all GCs in the V_321_L mice (**Figure 4b’’,** Mann-Whitney Test p=0.03, Two-stage step-up Benjamini-Krieger-Yekutieli FDR corrected for multiple comparisons q=0.002, U=0). Intriguingly, this increase in numbers of *marker*+*Penk*+ cells was only observed in *Penk* and *Nrg1* co-expressing GCs (**Figure S5A**) and not in *Nrg1* non-expressing *Penk*+ GCs (**Figure S5B**). In FISH experiments for *Penk* and *Nrg1*, we found that majority of the *Penk*+ GCs co-expressed *Nrg1* (∼94%) (**Figure S5B, inset**). We next performed FISH to detect co-expression of *Penk* with candidate markers like *Nptx2*, *Sorcs3,* and *Scg2* and found strong overlap between *Penk* and the selected marker genes, particularly biased towards the GCL-IML boundary of the suprapyramidal blade of the DG (**Figure 4C**). We selected *Nptx2* and *Sorcs3* given their higher relative exclusivity for expression towards the GCL-IML boundary compared to *Scg2*. We quantified the degree of overlap between *Penk* and *Nptx2* or *Sorcs3* and of all three genes using FISH. We found that on average 84% of the *Penk*+ GCs also expressed *Nptx2*, and 67% of the *Penk*+ GCs also expressed *Sorcs3*. Co-expression of *Nptx2* and *Sorcs3* was detected in 64% of the *Penk*+ GCs (**Figure 4D and 4d’**). Taken together with the restriction of *Penk* and *Sorcs3* co-expressing cells in the cell body layers at the GCL-IML boundary (**Figure 4a’**) and the recently reported SGC-like morphology of *Penk*+ GCs (Mortessagne, Cartier et al. 2024), these data indicate that the Penk, Nptx2, and Sorcs3 co-expressing population of GCs represent SGCs.

### Non-SGCs harbor the potential to express an SGC-like transcriptomic state

SGCs are thought to be generated exclusively during embryonic development, however, several lines of evidence suggest that the adult DG possesses the capacity to produce SGCs (Lauer, Beckmann et al. 2003, Kim, Duan et al. 2009, Fitzsimons, van Hooijdonk et al. 2013, Llorens-Martin, Jurado-Arjona et al. 2014, Terreros-Roncal, Flor-Garcia et al. 2019, Mortessagne, Cartier et al. 2024). We found that the SGC-enriched cluster of GCs (cl.18) was closely related to other GC clusters. Marker gene discovery was unable to identify any gene with unique expression that distinguished clusters 0, 1, and 5 from each other or from cluster 18 (**Figure S3D**). However, cluster 18 GCs showed expression of *Sorcs3* as a unique marker suggesting that the SGC-like transcriptomic state could be conceptualized as a modular “add-on extension” to a general GC-like transcriptome. To test this “add-on” concept, we performed trajectory analysis on the GC clusters using Monocle3 (Cao, Spielmann et al. 2019). To assign the root node, we assessed the expression of *Camk4*, which has been shown to transiently increase in expression in maturing GCs and *Ntng1*, whose expression gradually increases and peaks as GCs mature (Hochgerner, Zeisel et al. 2018) (**Figure 5A and 5a’**). FISH for *Camk4* and *Ntng1* revealed that they were expressed in a spatial gradient in the DG GCL such that *Camk4* expressing cells occupied the lower half of the GCL in proximity to the hilus (Hi), while the *Ntng1* expressing cells occupied the upper half in proximity to the molecular layer (MOL) (**Figure 5A**). Since the DG laminates “outside-in”, *Camk4*+ GCs represent GCs with a more recent birthdate compared to *Ntng1*+ GCs. We examined the expression of *Camk4* and *Ntng1* in our GC clusters and found opposing gradients of expression in the UMAPs such that cluster 0 GCs represented the younger GCs occupying the bottom half of the GCL, whereas GCs located along the continuum from cluster 0 to cluster 18 showed gradually increasing *Ntng1* expression, likely representing GCs located in the top half of the GCL (**Figure 5a’**). Next, we computed pseudotime for the cells falling along these trajectories (**Figure 5B**). The top gene explaining pseudotime in WT DG was found to be *Sorcs3*, the cluster 18 marker gene which we also found to be a marker gene for SGCs (**Table S2**). In line with this, cells with late pseudotime were found towards the end of cluster 1 and in cluster 18, where we previously noted presence of *Penk*+ GCs (**Figure 3D**). Trajectories in the V_321_L DG were highly altered and cells with late pseudotime were present in cluster 5 (**Figure 5B**, **right**). The cluster 1-18 transition area in V_321_L clusters fell along an intermediate pseudotime, however, *Sorcs3* expression still faithfully represented the cells in this zone indicating preserved molecular profile of SGCs in the V_321_L DG.

**Figure 5.**
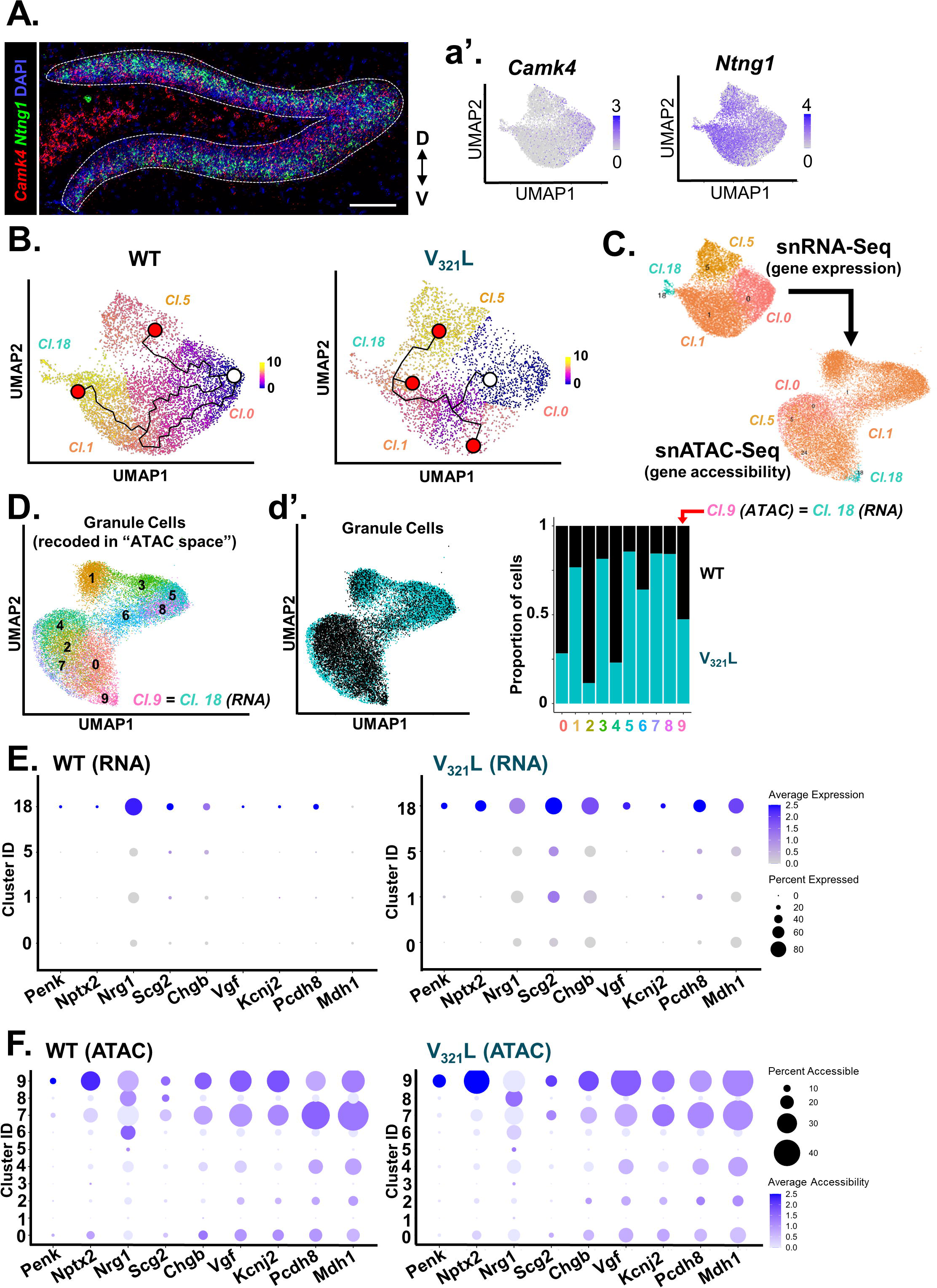
Non-SGCs harbor the potential to express an SGC-like transcriptomic state. **A.** Representative image of a WT DG showing expression of *Camk4* (red) and *Ntng1* (green). The DG blades are outlined with a white dashed line. Scale bar= 100µm. Anatomical legend indicates orientation of the slice (D= dorsal, V=ventral). **a’.** UMAPs showing expression of the early-stage maturation marker gene *Camk4* (left) and late-stage maturation marker *Ntng1* (right) with opposing gradients of expression within the granule cell clusters similar to their opposing gradients of expression within the GCL as shown in panel A. **B.** Trajectory mapping using Monocle3 reveals different trajectories between WT and V_321_L GCs. Pseudotemporal ordering of cells was performed by assigning root nodes in cluster 0 owing to higher *Camk4* expression. (**Left**) In WT GCs, Cluster 18 and adjoining cells in cluster 1 have the highest pseudotime, whereas cluster 5, which hosts one of the trajectory end points, is at an intermediate pseudotime. (**Right**) In V_321_L GCs, Cluster 18 and adjoining cells in cluster 1 have an intermediate pseudotime with a branched trajectory terminating at that junction. Meanwhile cluster 5 houses cells with the highest pseudotime. White circle indicates the root node for the trajectories and the red circles indicate end points for the different trajectories. **C.** GCs from snRNASeq clusters (clusters 0,1,5, and 18) were assessed for their chromatin accessibility using ATAC-Seq. UMAPS show GC clusters based on their gene expression profiles (RNA-Seq, **Top**) and the same GCs clustered based on their gene accessibility profiles (ATAC-Seq, **Bottom**). Colors and cluster IDs indicate identity based on gene expression. **D.** GCs were clustered based on gene accessibility (ATAC-Seq) and assigned new cluster IDs. Note that RNA-Seq cluster 18 GCs maintained their own cluster in the “ATAC space” as cluster 9. **d’.** (**Left**) UMAP of snATAC-Seq showing clusters of WT (black) and V_321_L (teal) nuclei. (**Right**) Quantification of proportional makeup of each of the snATAC-Seq clusters based on the genotype (WT vs. V_321_L) of the nuclei. Except for cluster 9, all the other clusters were either biased towards WT or V_321_L nuclei. **E.** Dot plot for 9 example SGC-enriched genes showing average gene expression (snRNA-Seq) represented by the color of the dots. Percentage of cells with expression of the specified gene within a given cluster is represented by the size of the dots according to the accompanying legend. Data from GCs – clusters 0,1,5, and 18 are shown split by genotype – WT (**Left**) and V_321_L (**Right**). Note that expression of SGC-enriched genes is largely restricted to cluster 18 in WT GCs. **F.** Dot plot for 9 example SGC-enriched genes showing average gene accessibility (snATAC-Seq) represented by the color of the dots. Percentage of cells with accessibility of the specified gene within a given cluster is represented by the size of the dots according to the accompanying legend. Data from GCs – clusters 0-9 are shown split by genotype – WT (**Left**) and V_321_L (**Right**). Note, cluster 18 GCs from Panel E are represented in cluster 9. Thus, genes enriched for expression in SGCs are broadly accessible to GCs.

Taken together, these data indicate that SGCs can be placed along a molecular continuum of GC age and that the SGC transcriptomic phenotype is largely unaltered in the V_321_L DG. Acquisition of an SGC-like transcriptomic state would necessitate accessibility of SGC-enriched genes along with presence of the appropriate transcriptional regulators which can activate/de-repress the SGC-enriched genes. To investigate the former of these two requirements, we examined chromatin accessibility in the same GCs that were profiled for gene expression. We remapped the GCs identified based on their gene expression profiles into the “ATAC space” (**Figure 5C**). We noticed that GCs from clusters 0 and 18 preserved their within cluster cohesion continuing to co-cluster in a single ATAC cluster each. However, GCs from clusters 1 and 5 were found to distribute into 8 smaller sub-clusters (**Figure 5D**). These results potentially indicate a more dynamic restructuring of the chromatin landscape in GCs that represented the intermediate pseudotimes. We also noted that there was a dramatic difference in the distribution and clustering of GCs from WT and V_321_L DG based on chromatin accessibility (**Figure 5d’, left**). Cluster 9 comprised of the *Penk*+ GCs and it was uniformly populated by WT and V_321_L GCs, however, cluster 4 which comprised of the *Camk4*+ GCs (cluster 0 in the “RNA space”) was disproportionately populated by WT GCs, which made up ∼75% of all cells in that cluster. These results indicate that V_321_L GCs might escape a “younger” chromatin state. To investigate this further, we performed trajectory analysis and pseudotime mapping of the ATAC-Seq data (**Figure S6**). Once again, we found that the *Penk*+ GCs (now located in ATAC cluster 9) had the maximum pseudotime values based on chromatin accessibility (**Figure S6A, left**). Intriguingly, in the analysis of WT GCs, there were three end points to the trajectories, one of which terminated at the cluster with the maximal pseudotime. However, in the V_321_L GCs, there was only one end point terminating at cluster 9 (**Figure S6A, right**). We evaluated the accessibility of the *Penk* transcription start site (TSS) (±1kb) and found that accessibility of the *Penk* TSS increased along pseudotime in both WT and V_321_L GCs, but the trajectories traversed different clusters (**Figures S6B and S6b’**). Surprisingly, unlike the WT-V_321_L difference in total number of cells expressing *Penk* (**Figure 1F**), the total number of GCs having the *Penk* locus accessible was not significantly different between the genotypes (**Figure S6b’’;** Mann-Whitney Test p=0.9 U=7). Therefore, next we compared the expression and accessibility of the SGC-enriched genes in WT and V_321_L GCs. We examined the expression of a select set of SGC-enriched genes for which there were RNA and ATAC counts present for each cell. As shown earlier, there were more cells expressing SGC-enriched genes in V_321_L mice and the expression was not as restricted to cluster 18 compared to the WT GCs (**Figure 3D**). Cluster 18 GCs (RNA-Seq) were remapped to cluster 9 (ATAC-Seq), and we found accessibility for each of the marker genes was present in cluster 9 of both WT and V_321_L GCs (**Figure 5F**). However, we also found SGC-enriched genes to be accessible in cells in nearly all other clusters for both genotypes.

These data indicate that GCs in WT mice might utilize an active repressive mechanism to prevent expression of SGC-marker genes in non-SGCs, whereas this mechanism might be disrupted in the V_321_L DG. To investigate this further, we searched for putative regulators for the SGC-enriched genes using ChEA and ENCODE databases. We identified 60 genes whose expression was enriched in Penk+ GCs and which were also correlated with maximal pseudotime in the trajectory analysis of GC gene expression (**Figure 5B; Table S2**). We found strong enrichment for regulation of these genes by Suz12, a core component of the polycomb repressive complex 2 (PRC2) (**Table S3**).

Thus, taken together these data indicate that an SGC-like transcriptomic state is accessible to non-SGCs, but is kept repressed by Nrg1 nuclear back signaling potentially through regulation of the PRC2 (Rajebhosale, Jone et al. 2024). Loss of Nrg1 nuclear back signaling due to V_321_L mutation might result in de-repression of the SGC genes and subsequent acquisition of an SGC-like transcriptome by non-SGCs.

### SGC numbers in the DG increase during the adolescence-adulthood transition in wild-type mice

Our data predicted that repression of SGC genes is mediated at least in part by PRC2. The role of PRC2 in lineage specification is well studied; recent reports have shown that PRC2 dictates the timeline of neuronal maturation (Ciceri, Baggiolini et al. 2024). In line with this, a recent study showed age-dependent increase in *Penk* expression in the DG GCL, which was not associated with adult-born or immature GCs (Mortessagne, Cartier et al. 2024). Thus, using co-expression of *Nptx2*, *Sorcs3*, and *Penk* as an SGC “identifier” we quantified SGCs in the mouse DG of 1-, 2-, 3-, and 12-month-old WT mice (**Figure 6A and 6B**). We confirmed that WT mice indeed show increase in *Penk* expression starting between 2-3 months of age, a period marking puberty and transition to early adulthood (**Figure 6a’**; Welch ANOVA p=0.0032 W=36.28(3.000,3.724). Dunett’s T3 corrected multiple comparisons: 1- vs. 3-month p=0.008, 1- vs. 12-month p=0.03, 2- vs. 3-month p=0.008). We found that the numbers of *Penk+Nptx2+Sorcs3+* GCs also increased with age (**Figure 6b’**; Kruskal-Wallis Test p=0.0012, KW=9.5. 1- vs 12-month Dunn’s corrected p=0.039). These dynamics of SGC abundance over adolescence were lost in the V_321_L DG (**Figure S7A**). There were no significant differences in the overall percentage of *Nptx2* or *Sorcs3* expressing *Penk*+ GCs at any of the ages indicating that this combination of markers stably reports this population of GCs over aging in both genotypes (**Figure S7B**). Since *Penk*+ GCs are embryonically generated and bulk of the GCs are not, we wondered if the SGC-like transcriptome might simply reflect cellular age and not necessarily mark SGCs. We examined ISH data from embryonic day 18.5 (E18.5) mouse DG (Allen Brain Atlas), when the newborn SGCs would be between 0 and 3 days of age, for expression of a subset of the SGC-enriched genes from our snRNASeq analysis (**Figure S7C**). We found cells expressing SGC-enriched genes such as *Penk, Scg2, Vgf,* and *Cit* in the dorsal border of the developing GCL at E18.5 indicating that marker gene expression is unlikely to be an artifact of cellular age (**Figure S7C**).

**Figure 6.**
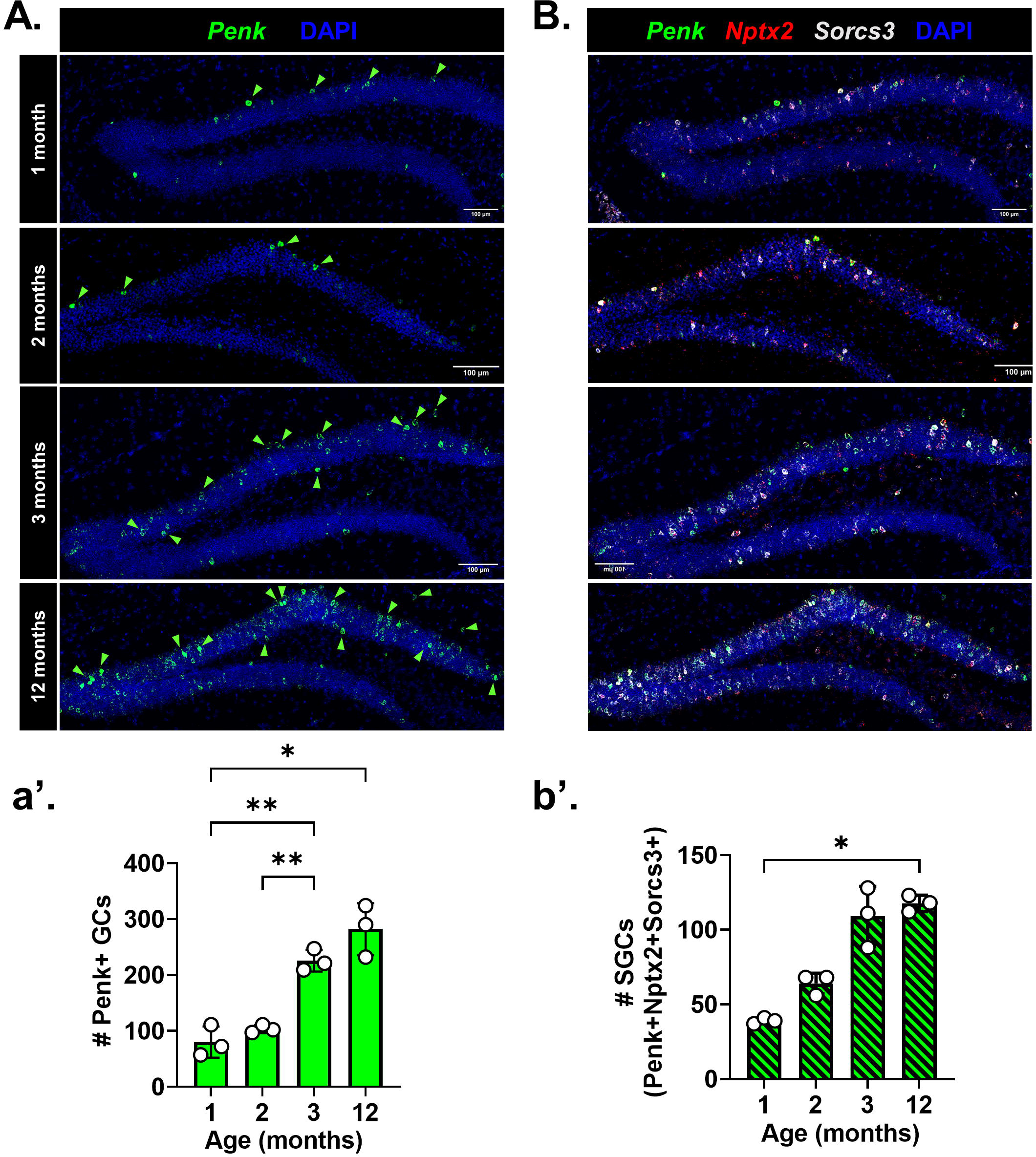
The total number of SGCs increases over the adolescence to adulthood transition in WT mice. **A.** Representative images of WT DG showing expression of *Penk* (green) using ISH at different ages: 1-, 2-, 3-, 12-months-old^$^. Green arrowheads indicate *Penk*+ cells in the GCL. Scale bar= 100µm. **a’.** Quantification of total number of *Penk*+ cells in the GCL of the suprapyramidal blade over age (Brown-Forsythe ANOVA p=0.002, F= 33.4(3, 4.2). Dunnett’s T3 multiple comparisons test: 1- vs. 3-months **p=0.008, 1- vs. 12-months *p=0.03, 2- vs. 3-months **p=0.008). ^$^Note that the 12 months old group comprised of animals aged 9-12 months of age. **B.** Representative images of WT DG showing expression of *Penk* (green), *Nptx2* (red), and *Sorcs3* (white) using ISH at different ages: 1-, 2-, 3-, 12-months-old^$^. Scale bar= 100µm. **b’.** Quantification of total number of *Penk*+*Nptx2*+*Sorcs3*+ cells in the GCL of the suprapyramidal blade over age (Kruskal-Wallis Test p=0.001, KW=9.5. Dunn’s multiple comparisons test: 1- vs. 12-months *p=0.039). ^$^Note that the 12 months old group comprised of animals aged 9-12 months of age.

Additionally, we did not find any *Penk*+ cells that expressed the immature neuron marker doublecortin (*Dcx*) at 3 months of age indicating that the source of the increase in SGC marker+ cells during the adolescence to adulthood transition is not likely to be immature GCs in line with predictions from the trajectory analyses which posit a gain of an SGC phenotype in mature GCs (**Figure S7D**). Thus, we sought to identify genes whose expression delineates the cell type transitions toward the SGC-enriched cluster 18 (**Figure S8A**). Using Monocle3 we identified modules of genes whose expression most strongly correlated with end-trajectories toward cluster 18 resulting in the identification of 4 significantly correlated modules (**Figure S8B**). Next we identified genes within these end-trajectory modules that overlapped with our previously identified SGC-enriched gene set with the rationale that genes whose expression delineates transitions toward cluster 18, and maintained higher expression in Penk+ GCs are more likely to represent key components of the SGC phenotype; we identified 60 genes which met this criteria (**Figure S8C**). Subjecting these genes to ontology analyses for cellular and biological processes, we identified them to be enriched for regulation of variety of neuronal functions with the feature of highest odds ratio being regulation of RMP, in line with our results showing a more hyperpolarized RMP in morphologically identified SGCs (**Figure 1J**). This enrichment was driven by two potassium channel genes-*Kcnk1* and *Kcnj2* (**Figure S8D**). Examination of expression of these genes showed that both had a bias for expression along the pseudotime continuum shown in Figure 5, with more consistent and higher expression toward cluster 18 (**Figure 8E**). Like our findings regarding SGCs, we noted higher and widespread expression of both *Kcnk1* and *Kcnj2* in the V_321_L GCs (**Figure 8F**). Thus, the identified 60 genes are good candidates for assessment of unique functional aspects of SGCs. These 60 genes were also enriched for significant associations with a variety of neuropsychiatric phenotypes and traits such as sensory sensitization, psychosis, schizophrenia, bipolar disorder, and neuroticism (**Table S4**).

### Natural decline in capacity for Nrg1 nuclear back signaling corresponds to the SGC fate

SGCs express high levels of *Nrg1* implying a role for Nrg1 signaling in SGC function (**Figure 4B**). We also found that a loss of Nrg1 nuclear back signaling due to the V_321_L mutation resulted more SGCs implying that intact Nrg1 nuclear signaling suppresses the SGC fate (**Figures 2-4**). To reconcile these findings, we postulated that Nrg1 expressed in WT SGCs might have a reduced capacity to engage in nuclear signaling. Nuclear back signaling by Nrg1 has been shown to be carried out by the Type III isoform of Nrg1 (Bao, Wolpowitz et al. 2003). Thus, we examined the expression of Type III Nrg1, in the DG over age hypothesizing that a reduction in expression of Type III Nrg1 might correspond to the adolescence to adulthood transition period when the SGC-marker+ cells increase. We found that Type III Nrg1 expression declines from 1-month to 12-months of age in WT mice, however the reduction between 1 and 3 months of age was not significant (**Figure 7A**; Kruskal-Wallis Test p=0.01 KW=6.25. 1- vs. 12-months Dunn’s corrected p=0.04).

**Figure 7.**
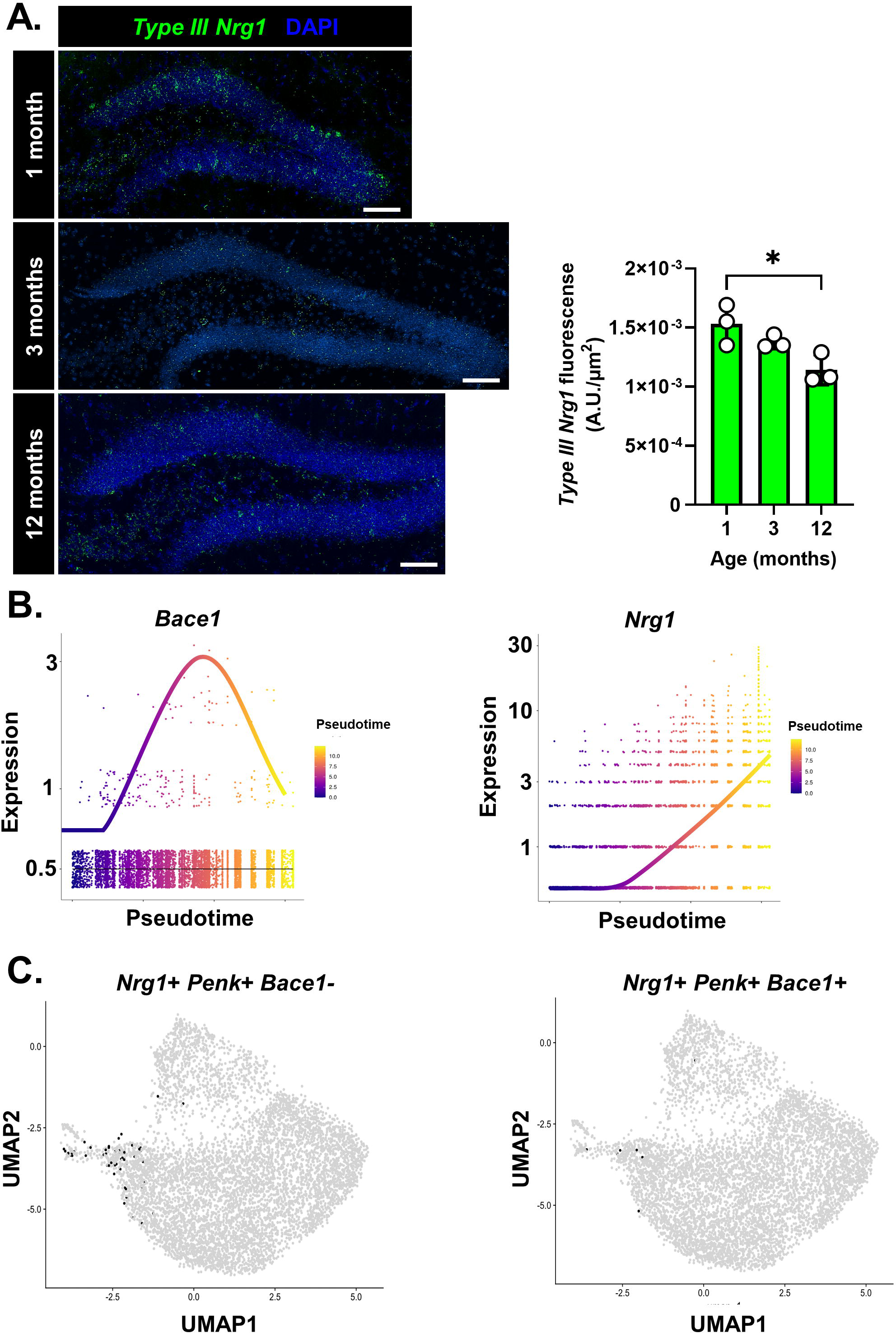
Age-dependent decrease in nuclear back signaling in the DG. **A.** (**Left**) *Type III Nrg1* expression declines with age in the DG GCL. Representative images showing FISH for *Type III Nrg1* (green) in WT DG over age (1-, 3-, and 12-months). (**Right**) Quantification of *Type III Nrg1* fluorescence density in the suprapyramidal blade of the DG shows a statistically significant decline in expression over age (Kruskal-Wallis Test p=0.01 KW=6.3. Dunn’s corrected multiple comparison: 1- vs. 12-months p=0.04). N=3 mice/age group. **B.** Pseudotime plots showing expression of *Bace1* (**Left**) and *Nrg1* (**Right**) in WT GCs. *Bace1* expression peaks at intermediate pseudotime and drops at maximal pseudotime, whereas *Nrg1* expression increases along pseudotime. **C.** UMAPs showing WT cells that express *Nrg1* and *Penk*, but not *Bace1* (**Left**) and cells co-expressing *Nrg1, Penk* and *Bace1* (**Right**). Each black dot represents 1 cell expressing the designated gene(s) and grey dots represent cells not expressing the gene(s).

*Bace1* is a critical regulator of nuclear back signaling. Cleavage by *Bace1* is necessary for producing Nrg1 proteins capable of being processed by gamma secretase (Willem, Garratt et al. 2006, Vullhorst, Ahmad et al. 2017). Thus, we examined expression of *Bace1* along pseudotime trajectories. Bace1 expression was found to be higher in cells at intermediate pseudotime compared to beginning and end of the trajectories, indicating low *Bace1* expression in SGCs (**Figure 7B**). Thus, while SGCs have higher expression of Nrg1 relative to non-SGCs (**Figure 7B**), they likely have low levels of nuclear back signaling due to lack of *Bace1*. Indeed, examination of cells expressing *Penk, Nrg1*, and *Bace1* in WT DG resulted in <10 cells showing co-expression of all three markers, while majority of the *Penk*+*Nrg1*+ GCs did not express detectable levels of *Bace1* transcripts (**Figure 7C**).

Thus, we conclude that the SGC-like transcriptomic state is actively repressed in the young DG by Nrg1 nuclear back signaling, and that some GCs acquire the SGC profile over adolescent development potentially due to a natural decline in Nrg1 nuclear back signaling.

## Discussion

Gargantuan bioinformatic efforts for cataloging brain cell types have produced hierarchical trees of hundreds of subtypes of classically recognized cell types, prompting many to ask how many bonafide cell types truly exist in the brain, how plastic or hardwired these cell types are, and more importantly, what bearing do they have on brain function? We suggest that a bottom-up understanding of cell type development would aid in answering some of these questions.

In this study we investigated subtype diversity of dentate gyrus granule cells and the potential relationship of subtype composition with disease-relevant genetic variation. We found that SGCs, a morphologically distinct subtype of GCs also showed distinct electrical properties (**Figure 1**). We identified unique genetic markers for SGCs and found an increase in the number of SGC marker expressing cells with age of the animal starting between 2 and 3 months of age (**Figures 4 & 6**). We assessed GC heterogeneity in a mouse model harboring a missense mutation in the *Nrg1* gene which was previously associated with psychosis (Walss-Bass, Liu et al. 2006). There were higher numbers of morpho-electric-transcriptomically defined SGCs in the mutant mouse DG even prior to 3 months of age (**Figure 2**). An analysis of GC chromatin accessibility revealed that SGC marker genes were accessible but not expressed in WT GCs indicating a repressive mechanism preventing access to an SGC-like transcriptomic state, and potential failure of this mechanism with disease-associated genetic variation (**Figure 5**).

### Composition of cell type identity: insights from the V_321_L DG

SGCs have unique morphological and intrinsic electrical properties (**Figure 1**). We found that all recorded neurons in the V_321_L DG, regardless of morphology, displayed SGC-like electrical properties (**Figure 2I-M**). We also noted that while there were no major morphological distinctions between WT and V_321_L GCs or SGCs, V_321_L GCs showed a wider dendritic splay than WT GCs, similar to that of SGCs (**Figure 2D**). These data indicate that the SGC fate might be driven by distinct modules of co-regulated genes involved in distinct cellular functions. Trajectory analysis revealed that SGCs were among cells with the maximal pseudotime (**Figure S6C and S6c’**). Gene Ontology (GO) analysis for genes whose expression was enriched in SGCs and those marking end trajectories in the WT DG showed “regulation of RMP” as the top function represented by *Kcnj2* and *Kcnk1* (**Figure S8D**). A more hyperpolarized RMP was one of the SGC-specific electrical properties (**Figure 1J**). In V_321_L mice, we found that GCs with a single primary dendrite also showed an RMP resembling that of SGCs (**Figure 2I**). Upon visualizing expression of *Kcnj2* in WT and V_321_L GC clusters, we observed that *Kcnj2* was typically (WT) expressed in cluster 18 and in cluster 1 along the pseudotime trajectory but not in clusters 0 or 5. However, in V_321_L GC clusters, *Kcnj2* expression was noted in clusters 0 and 5, which might indicate that these neurons represent the GCs with single primary dendrites and electrical properties of SGCs. A similar pattern was noted for *Kcnk1*.

Nrg1 back signaling operates through two known mechanisms-local activation of PI3K-Akt signaling at the membrane and nuclear signaling via translocation of the ICD from the membrane to the nucleus (Bao, Wolpowitz et al. 2003, Hancock, Canetta et al. 2008). The high expression of Nrg1 in SGCs with concomitant decrease in capacity for nuclear back signaling indicates a potential shift in the balance between local and nuclear back signaling mechanisms (**Figure 7**). Enhanced Akt signaling, either via constitutive activation or loss of negative regulators like Pten and Disc1, has been shown to result in the production of GCs with multiple primary dendrites (Kim, Duan et al. 2009). Thus, Nrg1 signaling might regulate the production of the SGC phenotype through gene regulatory and local signaling mechanisms to drive global changes in electrical properties and cytoskeletal dynamics.

We noted the presence of GCs with SGC-like electrical properties (**Figure 2**) in the V_321_L DG. This could potentially indicate a modular switch to SGC-like electrical properties while maintaining a typical GC morphology. Faster downstroke of the action potential was found to be one of the strongest cell type-defining features of SGCs (**Figure S2C**). However, downstroke was one of few electrical properties which retained correlation with GC position in the GCL even after exclusion of SGCs from the analysis indicating a relationship with GC birthdate (**Figure S1I**). We previously noted accelerated depletion and neuronal commitment of stem cells in the adult V_321_L DG (Rajebhosale, Jone et al. 2024). It is possible that this loss of regulated neurogenesis also occurs in the late embryonic and early postnatal stages resulting in greater production of the early-postnatal pool of GCs in V_321_L mice. Neonatally born GCs show some features of SGCs including wider splay angles as seen in the V_321_L DG (**Figure 2D**) (Save, Baude et al. 2019, Masachs, Charrier et al. 2021). Whether these features and the additional SGC-like cells in the V_321_L DG arise from accelerated early neurogenesis and/or altered specification of postnatal and adult neurogenesis awaits further inquiry.

### The Semilunar Granule Cell: cell type or an inevitable cell state?

We found that the SGC transcriptomic fate was accessible to non-SGCs and that the numbers of cells expressing SGC markers increased with age of the animal. However, we propose that this does not indicate the presence of a single GC type representing an inevitable fate for all GCs. In the WT DG, we found that the pseudotime analysis accurately captured chronological time, ordering cells based on their birthdates (**Figure 5A & 5B**). We found only one trajectory toward the *Penk*+ GCs (cluster 18) which traversed cluster 1. There was a separate trajectory ending in an intermediate pseudotime in cluster 5, with clear separation between clusters 5 and 18 (**Figure 5A & 5B**). Thus, we propose that the morphological distinction between GCs and SGCs faithfully represents two distinct types of GCs. However, there exists a third type of GC (clusters 0 and 1), which harbor the potential to acquire the SGC fate. This is apparent in the analysis of *Kcnj2* expression mentioned earlier-*Kcnj2* expression increases along pseudotime in cluster 1 but not in cluster 2 despite having identical pseudotime values. This can also be seen in the analysis of end-trajectory gene modules: clusters 1, and 18 have more common gene modules correlated with their identities whereas cluster 5 often shows anticorrelation in those modules (**Figure S8**). Our analyses indicate that the access to the SGC fate might be regulated by an “epigenetic” barrier deposited by the PRC2, and erasure of this barrier and a potential concomitant activation might underlie regulation of cell type composition in the DG (**Table S3**). We have previously shown that the V_321_L DG shows significantly reduced expression of *Ezh2,* which encodes the histone methyltransferase necessary for PRC2 function, and a concomitant increase in the expression of genes predicted to be PRC2 targets (Rajebhosale, Jone et al. 2024). Thus, the Nrg1 ICD might communicate developmental context such as synaptogenic interactions and/or synaptic activity to the genome through a PRC2 intermediary.

Given the unique circuit properties of SGCs and their potential to release enkephalin (encoded by *Penk*), we suggest that such regulation might afford the DG some flexibility to regulate cell type composition depending on the context of the circuit, to maintain proper circuit dynamics (Larimer and Strowbridge 2010, Gupta, Proddutur et al. 2020).

## Materials and Methods

### Experimental model and subject details

Male and female mice ages P24-12 months were used. Animals were housed in a 12-hour light/dark cycle environment that was both temperature and humidity controlled. Animals had free access to food and water. All animal care and experimental procedures were approved by the Animal Care and Use Committee of NINDS, National Institutes of Health, Bethesda MD (Animal protocol #1490).

### Genotyping

Genotypes were determined by PCR using the following primers:

Forward primer 5’- GGTGATCCCATACCCAAGACTCAG -3’

Reverse primer 5’- CTGCACATTTATAGSGCATTTATTTTG -3’

### Acute slice electrophysiology

Coronal brain slices containing hippocampus were prepared from P24-P34 days old mice. Animals were anesthetized with a mixture of ketamine and xylazine (100 mg ketamine and 6 mg xylazine/kg body weight injected ip). Then the brains were transcardially perfused with a sucrose-based solution. This solution contained (in mM): sucrose 230; KCl 2.5; MgSO_4_ 10; CaCl_2_ 0.5; NaH_2_PO_4_ 1.25; NaHCO_3_ 26; glucose 10 and pyruvate1.5. After decapitation, the brain was transferred quickly into the sucrose-based cutting solution bubbled with 95%O_2_ and 5% CO_2_ and maintained at ∼3°C. Coronal brain slices (300µm) were prepared using a Leica VT1200S vibratome (Leica, Inc). Slices were equilibrated with oxygenated artificial cerebrospinal fluid (aCSF) at room temperature (24-26°C) for at least 1 hour prior to transfer to the recording chamber.

Slices were moved to a stage of an upright, infrared-differential interference contrast microscope for patch clamp recording (SliceScope, Scientifica). The recording chamber was continually perfused with an artificial CSF at a rate of 2ml/min containing (in mM): NaCl 126, KCl 2.5, NaH_2_PO_4_ 1.25, NaHCO_3_ 26, CaCl_2_ 2, MgCl_2_ 2 and glucose 10 bubbled with 95% O_2_ and 5% CO_2_ to maintain pH at ∼7.4 with osmolarity of 305-315 mOsm at 31°C.

DG neurons were visualized with a 40 X water-immersion objective by infrared microscopy (Scientifica Pro, Scientifica). Patch electrodes with a resistance of 4–6 MΩ were pulled with a laser-based micropipette puller (P-2000, Sutter Instrument Company). Signals were recorded with a Multi Clamp 700B amplifier and pClamp11 software (Molecular Devices, Inc). The pipette solution contained (in mM) 125 K-gluconate, 10 KCl, 10 HEPES, 4 NaCl, 4 Mg-ATP and 0.3 Tris-GTP, and 7 Phosphocreatine (pH 7.3-7.4, osmolarity 290-295 mOsm). Granule cells in the middle of the granule cell layer (GCL) from the dorsal blade of the DG were recorded. In experiments focused on recording SGCs, recordings were made focused on the top cell body layer of the GCL.

Twenty features were extracted from the responses to current steps. (1) Membrane potential (mV), measured in the absence of a current injection. (2) Sag potential (mV), measured in response to a -60 pA step and equal to the difference between the steady-state potential and the minimum potential. (3) Input resistance (MΩ), measured by the response to a -20 pA step. (4) Membrane time constant τ (ms), measured by the relaxation to a -20 pA step. (5) Rheobase (pA), the minimum current step of 500 ms duration needed to elicit an action potential. (6) Spike threshold (mV), measured from the first action potential of the rheobase current step (“first action potential”) and defined as the potential at which dV/dt crosses 10 mV/ms. (7) Spike amplitude (mV), measured from the first action potential and defined as the difference between the tip of the action potential and spike threshold. (8) Spike width (ms), measured from the first action potential and defined as the width at half maximum (halfway between threshold and tip). (9) Spike latency (ms), measured at rheobase current and defined as the time difference between the start of the current step and the threshold crossing of the first step. (10) Spike peak (mV), the absolute value of the tip of the first action potential. (11) Upstroke (mV/ms), the maximum value of dV/dt on the upstroke of the first action potential. (12) Downstroke (mV/ms), the minimum value of dV/dt on the downstroke of the first action potential. (13) AHP amplitude (mV), measured from the aftermath of the first action potential and defined as the difference between threshold and the minimum potential within 100 ms after the action potential. (14) AHP latency (ms), the time between threshold crossing by the first action potential and the AHP minimum. (15) AHP width (ms), the time difference at half maximum of the first AHP. (16) f-I slope (Hz/pA), the slope of the initial linear section of the f-I curve. (17) Max firing rate (Hz), the maximum firing rate produced by a current step between 0 and 200 pA, across the entire 500 ms duration. (18) Adaptation index (dimensionless), measured from the maximal current step and defined as the number of spikes elicited in the second half of the step divided by the number elicited in the first half. (19) Coefficient of variation (CV) of interspike intervals, measured from the maximal current step. (20) AHP maximum (mV), measured from the maximal current step and defined as the difference between the baseline potential and minimum membrane potential recorded within 200 ms of the cessation of the step.

### Tissue processing

Animals were deeply anesthetized with isofluorane and transcardially perfused with 4% paraformaldehyde (PFA) in PBS. Brains were harvested and post-fixed overnight at 4°C in 4% PFA in PBS. Brains were transferred to 30% sucrose in PBS, incubated at 4°C with agitation until they sank and then embedded in optimal cutting temperature (O.C.T.) compound. After this, brains were flash frozen and stored at -80°C. 15μm sections were obtained serially from Bregma -1.5 to -2.5mm (approx). Sections were collected directly onto superfrost charged slides such that each slide contained 2 sections – one from a more anterior and one from a posterior hippocampal region. Slides were stored at -20°C in a box along with a desiccant.

### Immunohistochemistry

Following acute slice recordings, 300μm sections were fixed in 4% PFA at 4°C on a shaker overnight. The following day, sections were rinsed three times in 1xPBS and incubated in a fourth wash for 15 min at room temperature with constant agitation. Sections were then blocked with 10% normal donkey serum in 1xPBS containing 0.5% TritonX-100 at 4°C overnight on a shaker followed by incubation with fluorescently conjugated streptavidin at 4°C overnight with constant agitation. Sections were washed three times in 1xPBS containing 0.1% TritonX-100 (PBS-T) and mounted on superfrost plus glass slides with mounting medium containing DAPI.

### In situ hybridization

In situ hybridization was performed using the RNAScope multiplex v2 assay (ACD) or RNAScope HiPlex assay (ACD) using manufacturer recommended protocols available at -https://acdbio.com/sites/default/files/UM%20323100%20Multiplex%20Fluorescent%20v2_RevB.pdf

Details regrading probes used can be found in the KRT.

### Imaging & image analysis

Imaging was done using a Nikon Ti2 spinning disk confocal microscope. All slides in an experiment were imaged using identical settings. Images were acquired by capturing the whole DG and using z-steps of 1µm using a 40x silicon oil immersion objective. Image analysis was performed in imageJ (FIJI). Images were z-projected using maximum intensity projection and regions of interest were manually outlined. Fluorescence intensity density was measured as average intensity normalized to the area of the superior blade of the DG. The cell counter plug-in was used to manually count cells expressing *Penk* and other marker genes. We counted cells expressing 5 or more small puncta of any given gene or those with large clusters of RNA molecules. For most of the probes used, the RNA expression profile was “cell filling”. To analyze cell positions, we measured the distance of individual cells to the GCL-IML boundary by measuring the length of a line drawn to be perpendicular to the GCL-IML boundary.

Imaged neurobiotin filled neurons were manually reconstructed and analyzed using the filaments feature in Imaris (Oxford Instruments). Splay angles were calculated in FIJI using the “Angle tool” by sequentially placing one point on the most laterally displaced dendritic termination, followed by one in the center of the soma and then on the opposite most laterally displaced dendritic termination. Cell position was recorded by finding the optical section containing a section through the center of the cell body. Vertical distance from the center of the cell body to the GCL-IML boundary was measured, manually considering the angle of the blade of the DG.

### Single-nucleus RNA+ATAC Seq

At approx. 10 weeks of age, 4 WT and 4 V_321_L mice (2M/2F) were deeply anesthetized using isoflurane and perfused with ice-cold 1xPBS in an ice trough. All surfaces and tools in direct contact with tissue were thoroughly cleaned with RNAseZap (Invitrogen) prior to onset of procedure. The brain was rapidly removed and sliced using a prechilled brain matrix. Brain slices were transferred to ice-cold PBS and the dentate gyrus was dissected out and flash frozen. Tissue was stored at -80 C overnight. The following day, single nucleus suspension was prepared from the frozen DG tissue using the Chromium nuclei isolation kit (10xGenomics; Cat#1000494) following manufacturer guidelines. Nuclei were counted and diluted appropriately to obtain a final concentration of ∼1000 nuclei/µL. Approximately 3000 nuclei per animal were subjected to the transposition reaction and transposed nuclei were loaded onto the 10x Chromium platform using the Next Gem Chip J (10xGenomics; Cat#1000230) for gem generation. ATAC and RNA-Seq libraries were prepared using components of the Chromium Next GEM Single Cell Multiome ATAC + Gene Expression Reagent Bundle (10xGenomics; Cat#1000283) according to manufacturer guidelines. Libraries were sequenced on an Illumina NextSeq550 (High throughput kit) with ∼25,000 reads per nucleus for RNA-Seq and ∼20,000 read pairs per nucleus for ATAC-Seq. The cellranger-arc-2.0.2 pipeline (10xGenomics) was then applied to the sequence output producing data files for Gene Expression and ATAC-Seq analysis.

### Gene Expression Analysis

Sequenced libraries were analyzed in R using functions supported in the “Seurat” package unless otherwise described (Hao, Stuart et al. 2023). Cells were filtered to keep only those that had: 1) a minimum number of detected counts > 500, 2) a maximum number of detected counts < observed mean + 2SD = 8916.1, 3) a minimum number of detected features > 250, 4) a maximum number of detected features < observed mean + 2SD = 3307.53, 5) a percent mitochondrial counts < observed mean + 1SD = 2.79%. Doublet filtering was performed using “scDblFinder” package. Cells observed to have a doublet score > respective library mean + 2SD were then removed. Data was normalized using the “Seurat::NormalizeData” function. Before principal components analysis, the top 3000 highly variable genes were selected and the “ScaleData” function was applied excluding mitochondrial genes. To determine the number of components to use in the clustering of cells, the “RunPCA” (npcs=200) and “ElbowPlot” (ndims) functions were used and the “RunPCA” function re-applied after using the number of components decided upon (npcs=75). “Harmony” was used to integrate libraries across datasets. For clustering of cells, the “FindNeighbors” (reduction=“harmony”, dims=1:75) and “FindClusters” functions were used over a range of resolutions from 0.04 to 3.5 in steps of 0.01. Cluster results were summarized using the “clustree” package and 1.8 selected as the optimal resolution. At this resolution, several clusters were manually merged then the “findDoubletClusters” function from the “scDblFinder” package used to screen for and remove doublet enriched clusters. The “computeDoubletDensity” function was then re-applied but this time without splitting the cells by library. Cells observed to have a doublet score > cross cell mean + 2SD were then filter removed and the “FindNeighbors” and “FindClusters” functions re-applied under the same conditions described. Cluster results were again summarized using the “clustree” package and 2.1 selected as the optimal resolution. At this resolution, several clusters were again manually merged then cells for each cluster z-scored using UMAP dimension 1 values and separately using UMAP dimension 2 values. Cells in z-score space were then plot inspected by cluster using the “DimPlot” function and thresholds defined and applied to filter remove cells having a z-score greater than the defined thresholds. To identify conserved markers across library class (WT and V_321_L), the “FindConservedMarkers” function was used. To identify nonconserved markers per cluster, the “FindMarkers” function was used. To identify differential features between library class within cluster, the “FindMarkers” function was used setting the “ident” parameters to “WT” and “V_321_L” respectively. Granule cell clusters were next subset and trajectory fit using the “Monocle3” workflow (Trapnell, Cacchiarelli et al. 2014, Qiu, Mao et al. 2017, Cao, Spielmann et al. 2019). Root selection was accomplished by inspecting the expression for “Camk4” and “Ntng1” (Hochgerner, Zeisel et al. 2018). The “graph_test” function (neighbor_graph=“principal_graph”) was used to identify features that explain pseudotime (q_value<0.05). To identify depleted vs enriched modules of features that explain pseudotime, the “find_gene_modules” function was used.

### ATAC-Seq Analysis

The “atac_peaks.bed”, “barcode_metrics.csv”, and “atac_fragments.tsv.gz” files produced for the “WT” and “V_321_L” Dentate gyrus samples were imported into R (https://cran.r-project.org/) using the “read.table” function. Functions available in the “Signac” and “Seurat” packages were interchangabley used for analysis. To create a working fragment object per sample, the “CreateFragmentObject” function was used. Cells with an “atac_fragments” value <= 500 were discarded with surviving cells used as input into the “FeatureMatrix”, “CreateChromatinAssay”, and “CreateSeuratObject” functions, producing one “Seurat” object per sample. For each of these objects, genomic annotations for “EnsDb.Mmusculus.v79” were added and quality metrics generated. To add the annotations, the “GetGRangesFromEnsDb” and “Annotation” functions were used. To generate the quality metrics, the “FractionCountsInRegion”, “NucleosomeSignal”, and “TSSEnrichment” functions were used. Metrics produced from these functions, along with those for “peak_region_fragments”, “nCount”, “pct_reads_in_peaks”, and “TSS_fragments” were used to filter remove cells from each object via the “subset” function. Specifically, cells that did not have a value within 2SD of the observed mean for each metric were removed. Post filtering, all “Seurat” objects were combined into one using the “merge” function and a unified set of peaks generated using the “reduce” function. This unified peak set was then filtered to keep only those with a “peakwidths” value < 10000 and > 20. The filtered peak set was then used to regenerate the individual “Seurat” objects as described and those objects combined into one object. The “RunTFIDF”, “FindTopFeatures”, “RunSVD”, “RunUMAP”, and “FindNeighbor” functions were then applied to the combined object (dims = 2:20, reduction = ‘lsi’) in preparation for clustering. For clustering, the “FindClusters” function was used over a range of resolutions from 0.04 to 3.5 in steps of 0.01. Cluster results were summarized using the “clustree” package and 1.5 selected as the optimal resolution. At this resolution, several clusters were manually collapsed then cells for each cluster z-scored using UMAP1 dimension values and separately using UMAP2 dimension values. Cells in z-score space were then plot inspected by cluster using the “DimPlot” function and filter thresholds defined. Cells having a z-score greater than the thresholds were removed. For surivivng cells, annotations were transferred from the Single-nucleus RNA-Seq Analysis and log normalized activities for each known gene calculated. For annotation transfer, the “FindVariableFeatures” function (nfeatures = 5000) was used in conjunction with the “TransferData” function (reduction = ‘cca’, weight.reduction = ‘lsi’, dims = 2:20). To calculate the log normalized gene activities, and to add them to the combined “Seurat” object, the “GeneActivity”, “CreateAssayObject”, and “NormalizeData” functions were used. Specific clusters representing granule cells were then subset based on annotation from RNASeq data, and the functions used prior, per preparation for clustering, clustering, and post clustering, used again. For this subset, 0.5 was selected as the optimal resolution. Differential peaks between clusters produced at this resolution were tested for using the “FindMarkers” function (test.use = ‘LR’). Overrepresented motifs available in the “JASPAR2020” database (collection = ‘CORE’, tax_group = ‘vertebrates’) were also tested for using the “FindMotifs” function. Post testing, results were annotated with the closest occuring known gene using the “ClosestFeature” function. In addition to this testing, trajectory fitting was performed using the “Monocle3” workflow. This workflow was applied in three separate tacts: 1) using all cells representing “WT” and “V_321_L” samples combined, 2) using cells for “WT” samples alone, 3) using cells for “ V_321_L “ samples alone. Root selection for each of these tacts was accomplished by inspecting the gene activities for “Camk4”, “Igfbpl1”, “Fxyd7”, “Dcx”, “Calb2”, “Calb1”, “Ntng1”, “Penk”, and “Sorcs2” using the “plot_cells” function. Post fitting, the “cor.test” function was used in conjunction with the “p.adjust” function (method = ‘BH’) to test for and identify genes having pseudotime correlated with gene activity (q_value < 0.05). Enriched terms for these genes were subsequently identified using the “enrichR” package (dbs = ‘GO_Molecular_Function_2023’, ‘GO_Cellular_Component_2023’, ‘GO_Biological_Process_2023’).

### Statistical analysis

Statistical analysis was performed using Prism (GraphPad). Normality was assessed using Shapiro-Wilkins and Kolmogorov-Smirnov tests. If data failed normality test, non-parametric stats were used. p-values were corrected for multiple comparisons as necessary using Bonferroni (parametric) and Dunn’s (non-parametric) post-hoc tests.

## Key Resources Table

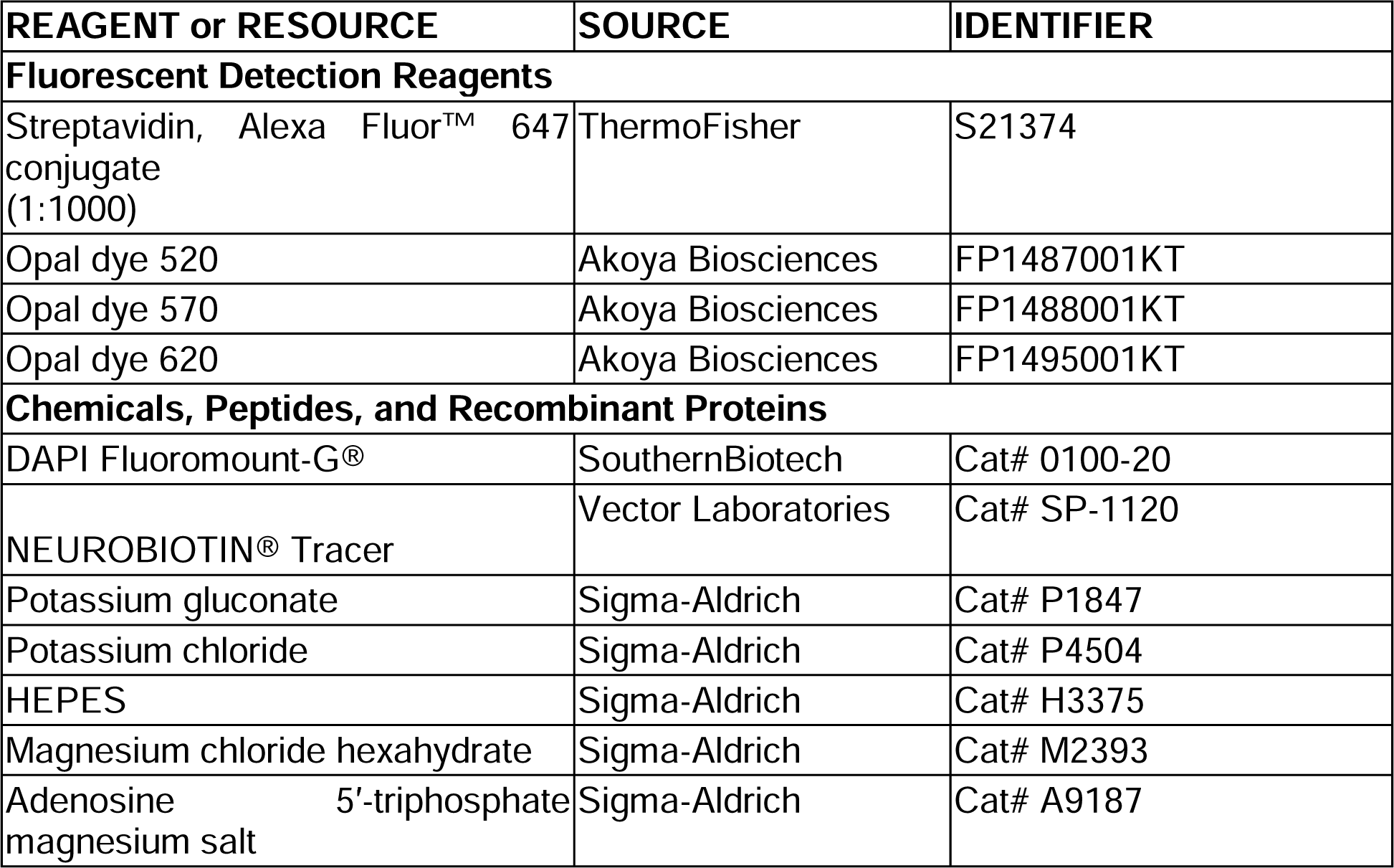
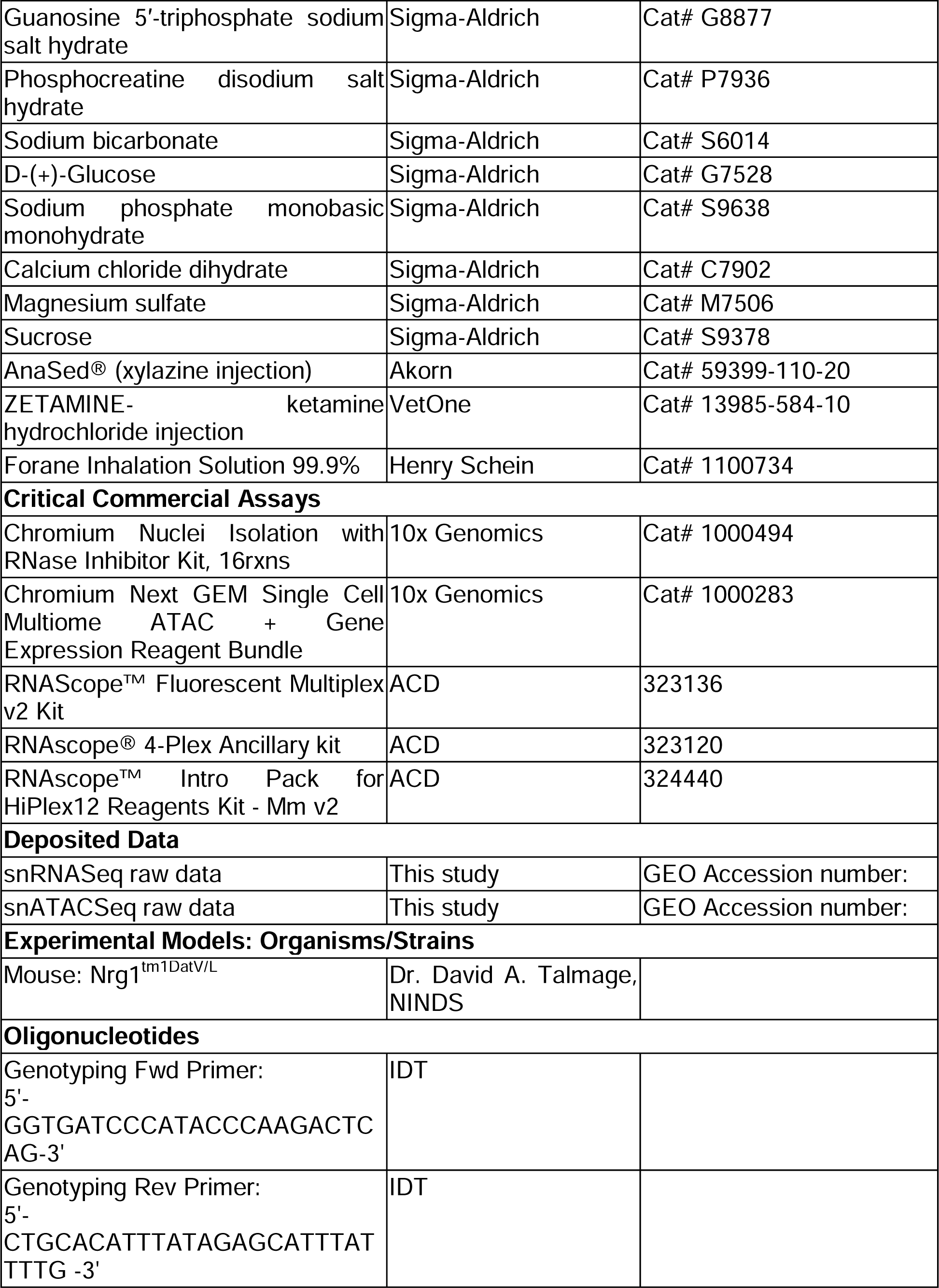
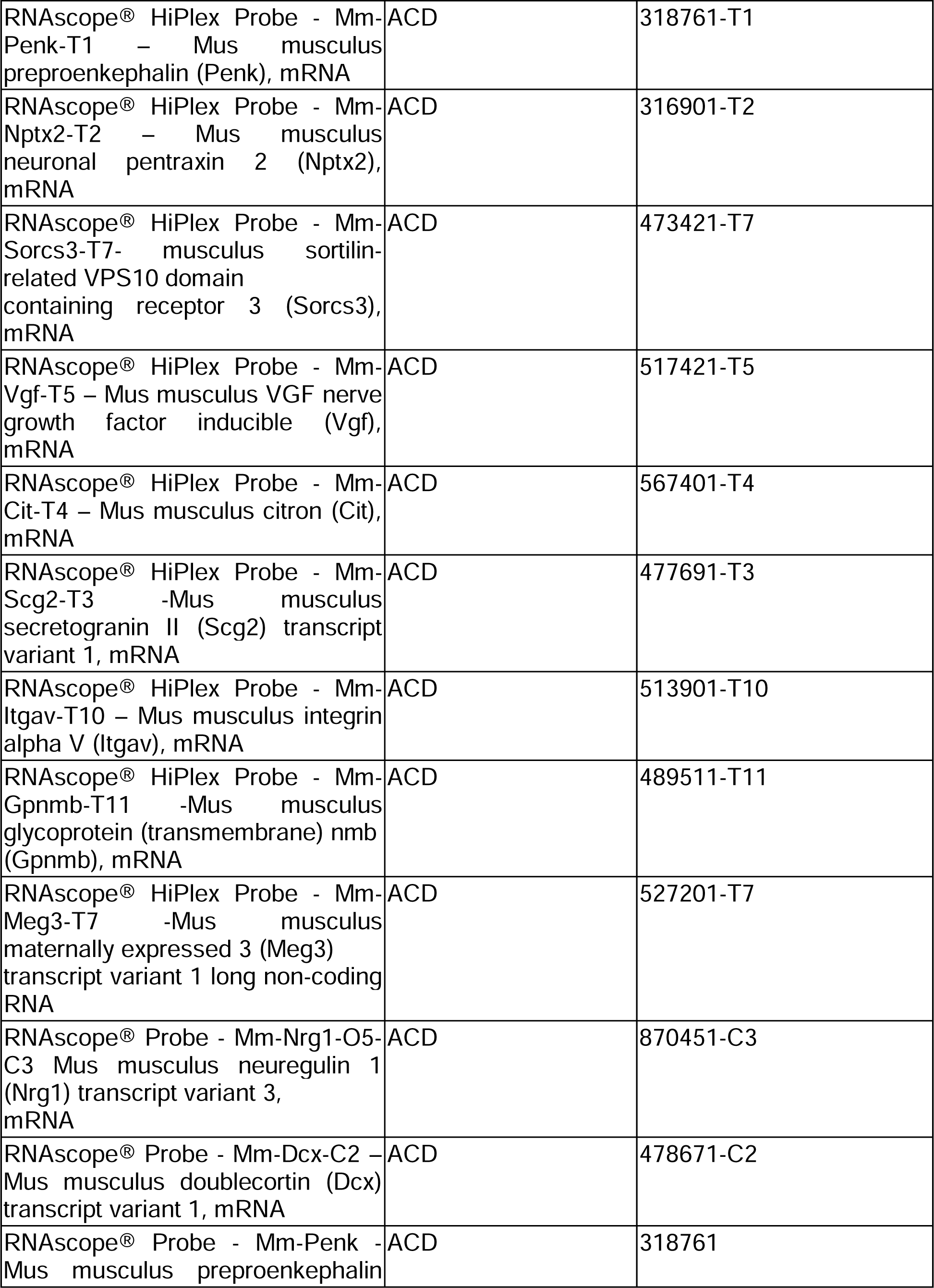
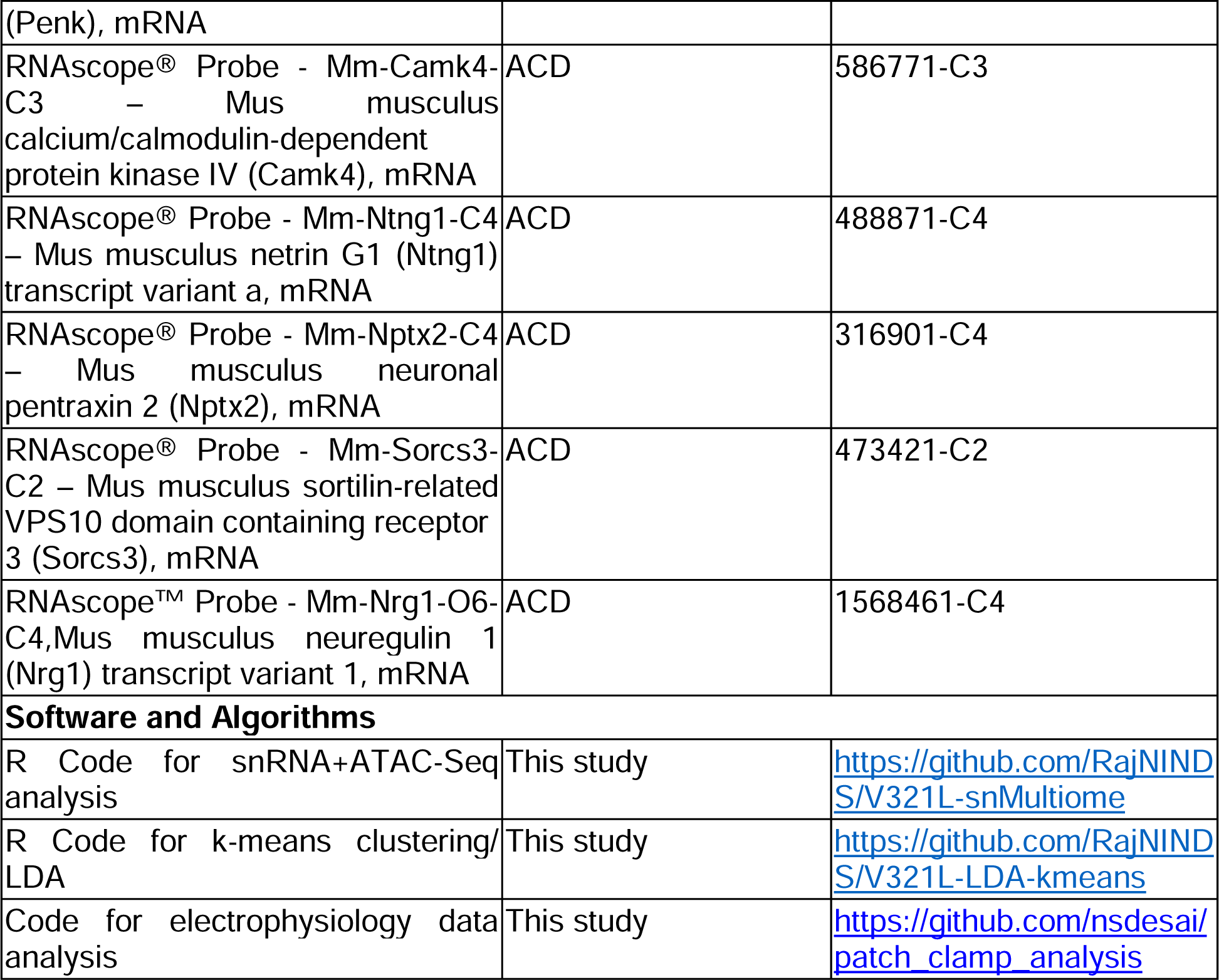

## Resource availability

### Lead Contact

Further information and requests for resources and reagents should be directed to and will be fulfilled by Lead Contact, Dr. David Talmage.

### Data and Code Availability

The data generated in this publication have been deposited in NCBI’s Gene Expression Omnibus (Edgar, Domrachev et al. 2002) and are accessible through GEO Series accession numbers provided in the KRT.

Code for data analysis can be accessed on Github using the links provided in the KRT.

## Acknowledgements

This work was supported by Intramural Research Program of NINDS. We would like to thank Dr. Abdel Elkahloun and Bayu Sisay (NHGRI sequencing core facility) for providing NGS consultation and services. In additional we would like to thank Taylor Muir for mouse colony management and Li Bai, Jessie Wallace, Shaina Sindhu, and Becca Alvarez for technical assistance with project management.

---

**Figure S1.**
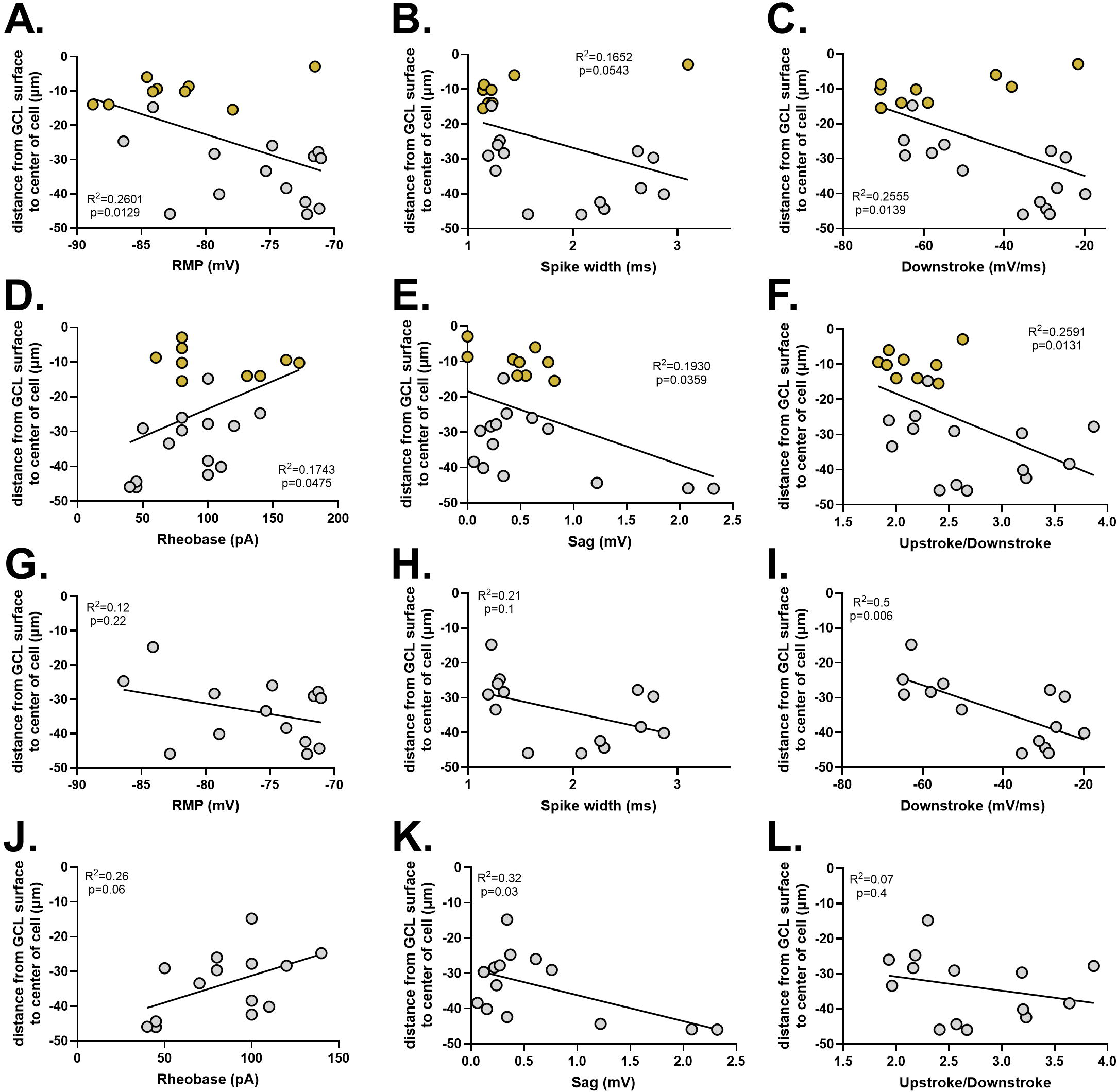
RMP and spike width are cell-type specific properties of SGCs. Relationship between neuronal cell body position and electrophysiological properties was evaluated. The following properties significantly co-varied with soma position: **A.** RMP (p=0.013, R^2^=0.26), **B.** Spike width (p=0.05, R^2^=0.17), **C.** Downstroke (p=0.014, R^2^=0.26), **D.** Rheobase (p=0.048, R^2^=0.17), **E.** Sag (p=0.036, R^2^=0.19), **F.** Upstroke/Downstroke (p=0.013, R^2^=0.26). N= 14 cells (GCs) and 9 cells (SGCs) from 5 mice. GCs are represented by silver circles and SGCs are represented by gold circles. Relationship between neuronal cell body position and electrophysiological properties was reevaluated for the above-mentioned properties while <u>excluding SGCs</u>: **G.** RMP (p=0.22, R^2^=0.12), **H.** Spike width (p=0.1, R^2^=0.21), **I.** Downstroke (p=0.006, R^2^=0.5), **J.** Rheobase (p=0.06, R^2^=0.26), **K.** Sag (p=0.03, R^2^=0.32), **L.** Upstroke/Downstroke (p=0.4, R^2^=0.07). N= 14 cells (GCs) from 5 mice.

**Figure S2.**
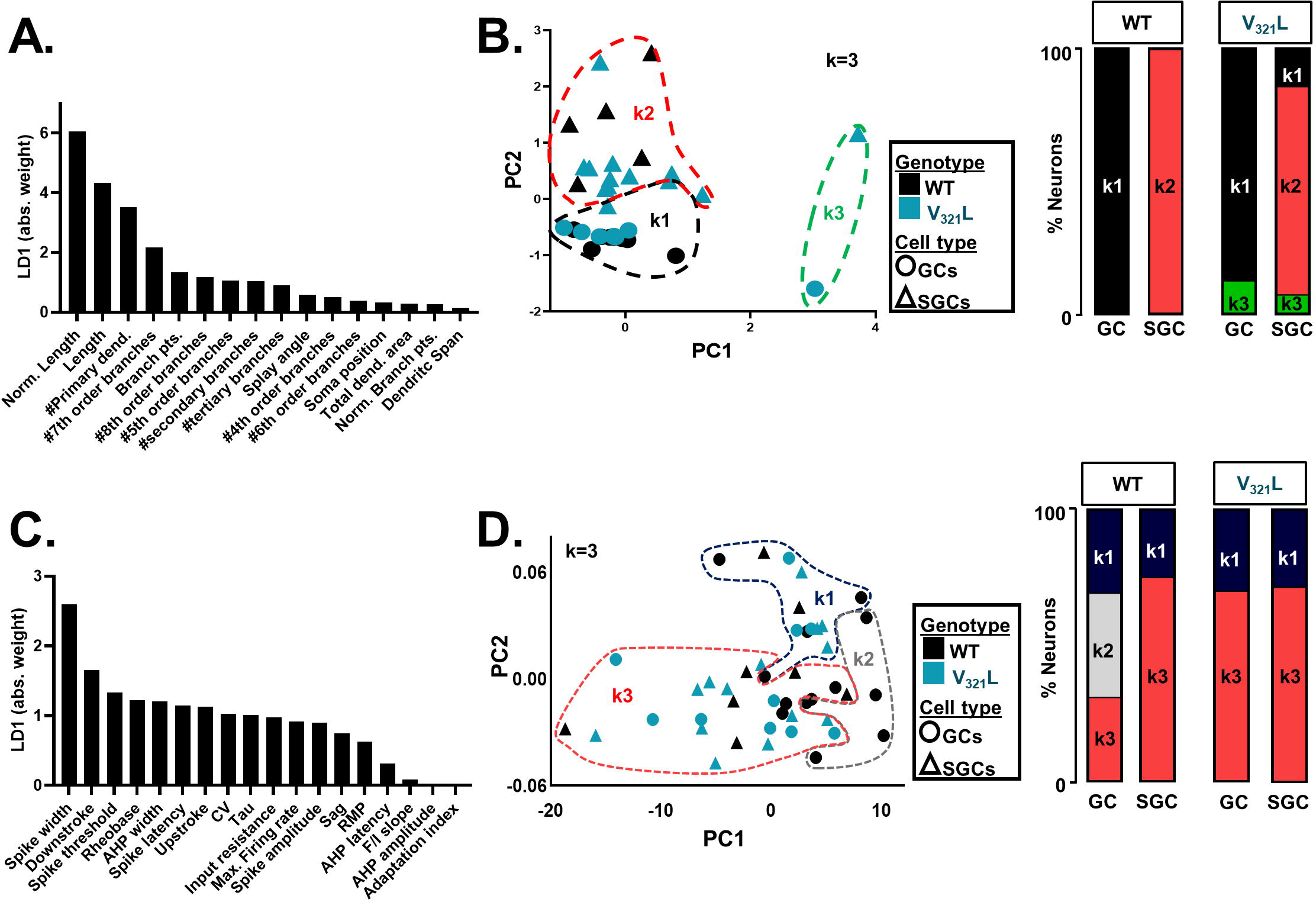
Supervised and unsupervised clustering of GCs and SGCs validate number of primary dendrites and spike width as important classifiers for SGCs. **A.** Plot showing contributions of individual morphological variables to LD1. Features strongly contributing to LD1 are dendritic length, number of primary dendrites, and branching. **B.** (**Left**) PCA plot. Each data point represents a single cell. Circles represent GCs and triangles represent SGCs. Black data points represent cells from WT mice and teal represents cells from V_321_L mice. k-means clustering was performed on morphological data with k=3. Dashed lines represent groupings of the three clusters – k1= black, k2= red, and k3= green. (**Right**) Proportional makeup of all GCs and SGCs from WT and V321L mice in relation to the k-means cluster they occupy. All WT GCs and SGCs segregate into k1 and k2 respectively. All but one V_321_L GCs occupy k1, while the one was in k3. All but 2 V_321_L SGCs occupy k2 with one in k1 and another in k3. **C.** Plot showing contributions of individual electrophysiological variables to LD1. Features strongly contributing to LD1 are spike width, and downstroke. **D.** (**Left**) PCA plot. Each data point represents a single cell. Circles represent GCs and triangles represent SGCs. Black data points represent cells from WT mice and teal represents cells from V_321_L mice. k-means clustering was performed on electrophysiological data with k=3. Dashed lines represent groupings of the three clusters – k1= navy blue, k2= grey, and k3= red. (**Right**) Proportional makeup of all GCs and SGCs from WT and V_321_L mice in relation to the k-means cluster they occupy. WT GCs are evenly distributed across all clusters whereas SGCs largely occupy k3 with a minority in k1. All V_321_L GCs and SGC cluster occupancy was virtually identical to WT SGCs.

**Figure S3.**
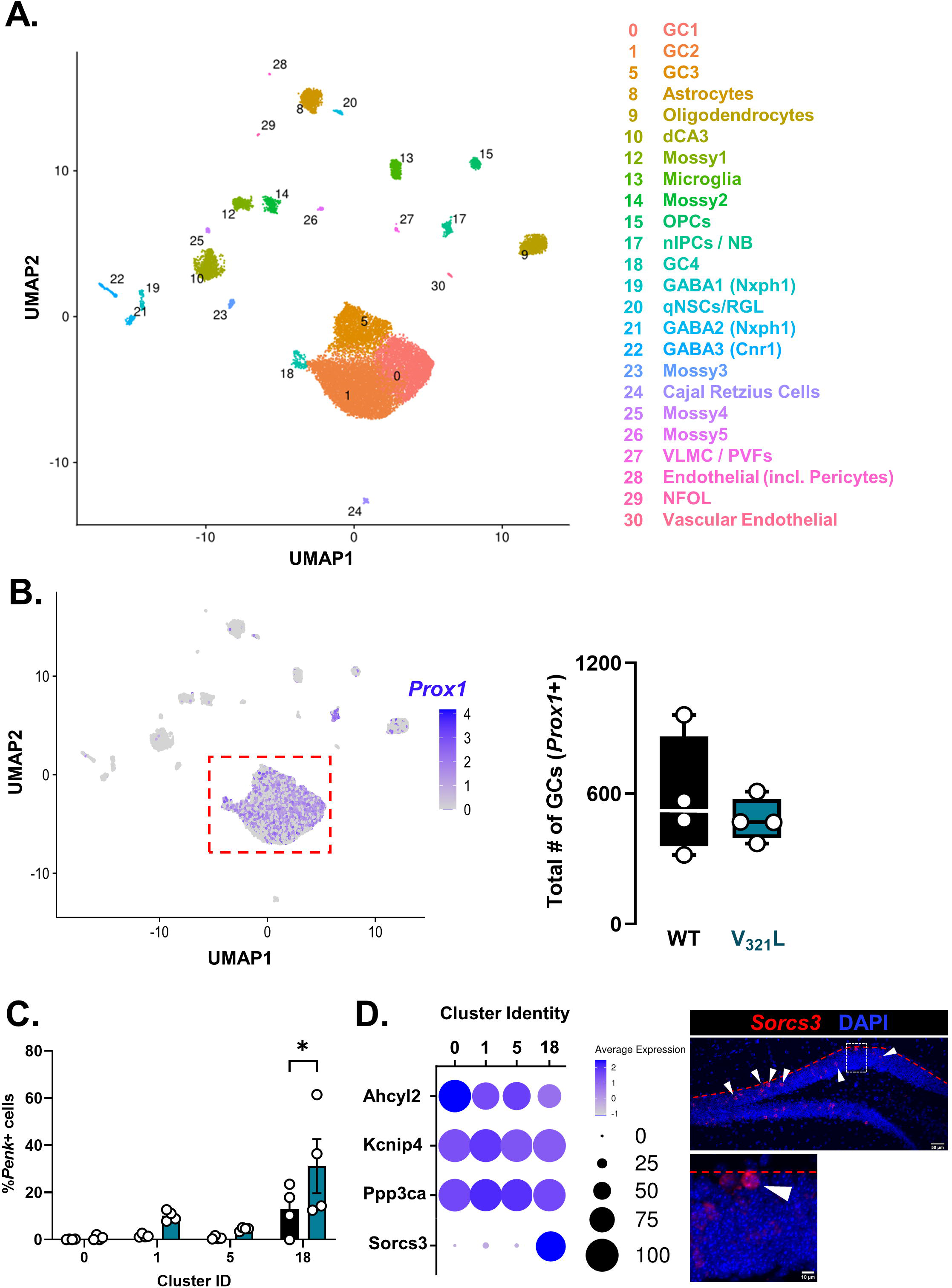
Expression of *Sorcs3* distinguishes SGC-enriched GC clusters. **A.** UMAP showing the analyzed snRNASeq data. Legend indicates cluster numbers and cluster identities. N=4 mice/genotype. Also see **Figure S4**. **B.** (**Left**) UMAP showing expression of the GC marker gene *Prox1* in all sequenced nuclei. (**Right**) Quantification of number of cells expressing *Prox1* from the snRNASeq GC clusters. There were no significant differences between WT (black) and V_321_L (teal) mice in the total number of *Prox1*+ cells (Welch’s t-test (two-tailed) p=0.5, t=0.6994, df=3.762). N=4 mice/genotype. **C.** Quantification of percentage of GCs expressing *Penk*+ in WT and V_321_L mice in individual GC clusters indicating cluster 18 as a cluster enriched for *Penk*+ GCs (SGCs). V_321_L mice had significantly higher enrichment of *Penk*+ GCs in cluster 18 compared to other GC clusters (Two-way RM ANOVA Cluster x Genotype p=0.2, Cluster p=0.0005, and Genotype p=0.054. Bonferroni corrected multiple comparisons: Cluster 0 vs. 18 ***p=0.0007, Cluster 1 vs. 18 *p=0.02, and Cluster 5 vs. 18 **p=0.003). N=4 mice/genotype. **D.** (**Left**) Results of conserved marker gene testing across samples for GC clusters. Dot plot shows expression of *Sorcs3* as the distinguishing feature of cluster 18. Legend alongside the dot plot indicates the color scale for average expression and size scale for proportion of cells expressing marker gene in each cluster. (**Right**) Representative image of a WT DG showing *Sorcs3* ISH (red) in the GCL. Inset shows a magnified image of an example cell expressing *Sorcs3* (arrowheads indicate *Sorcs3*+ cells in the GCL). Majority of the Sorcs3+ cells were found spatially biased towards the GCL-IML boundary (red dashed line) indicating that these cells are likely of embryonic origin.

**Figure S4.**
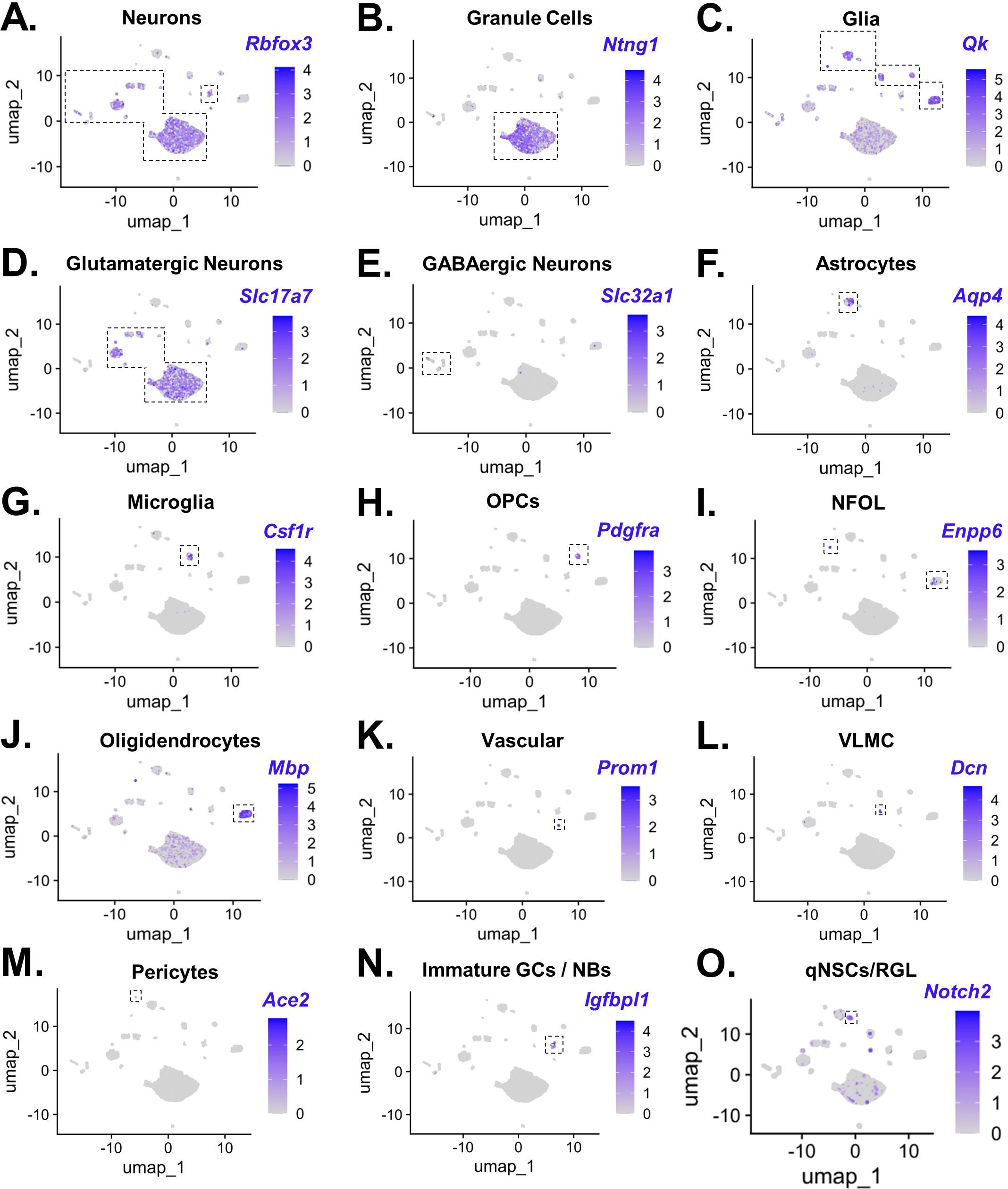
Cell type annotation of the snRNASeq data using marker gene expression. The following markers were used to identify cell types and annotate clusters-**A.** Rbfox3: pan-neuronal marker, **B.** Ntng1: granule cell marker, **C.** Qk: pan-glial marker, **D.** Slc17a7: glutamatergic neuronal marker, **E.** Slc32a1: GABAergic neuronal marker, **F.** Aqp4: astrocytic marker, **G.** Csf1r: microglial marker, **H.** Pdgfra: OPC marker, **I.** Enpp6: marker for newly formed oligodendrocytes (NFOL), **J.** Mbp: oligodendroglial marker, **K.** Prom1: vascular cell marker, **L.** Dcn: vascular leptomeningeal cell marker, **M.** Ace2: pericyte marker, **N.** Igfbpl1: immature GC/neuroblast marker, **O.** Notch2: marker for quiescent neural stem cells / radial glia-like cells.

**Figure S5.**
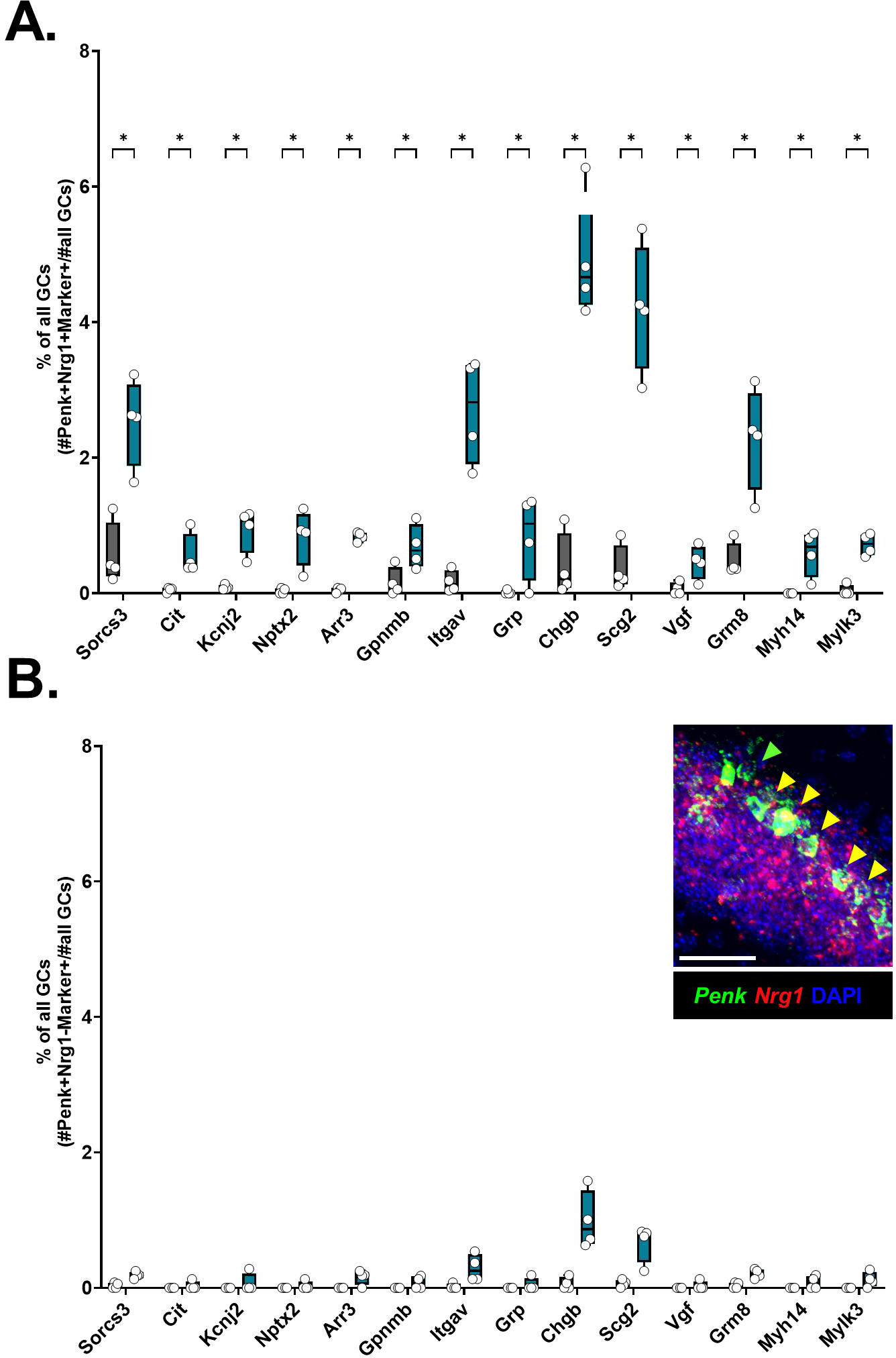
Increase in number of cells expressing SGC-enriched genes in V_321_L mice is specific to Nrg1-expressing SGCs. **A.** Quantification of precentage of *Penk*+*Nrg1*+ GCs expressing selected marker genes in WT and V_321_L mice. Number of cells co-expressing *Penk*, *Nrg1* and each of the marker genes were obtained from the snRNASeq data and divided by the total number of cells in clusters 0,1,5,18. WT and V_321_L samples were compared for each gene using multiple Mann-Whitney tests with an FDR correction. With the exception of Grp (WT vs. V_321_L FDR adj.p=0.14, U=2.5), V_321_L samples had significantly more cells co-expressing *Penk* and selected marker genes (FDR adj.p<0.1, U=0 or 1) (N=4 mice/genotype). **B.** Quantification of percentage of *Penk*+*Nrg1*-GCs expressing selected marker genes in WT and V_321_L mice. Number of non-*Nrg1* expressing cells co-expressing *Penk* and each of the marker genes were obtained from the snRNASeq data and divided by the total number of cells in clusters 0,1,5,18. WT and V_321_L samples were compared for each gene using multiple Mann-Whitney tests with an FDR correction. *Sorcs3*, *Itgav*, *Chgb*, *Scg2*, and *Grm8* (WT vs. V_321_L FDR adj.p=0.08, U=0), all other marker genes (WT vs. V_321_L FDR adj.p>0.1). Overall, the % of *Penk*+ GCs co-expressing marker genes was lower in *Nrg1* non-expressing GCs than *Nrg1*+*Penk*+ GCs (N=4 mice/genotype). (**Inset**) Representative image showing GCs expressing *Penk* (green), *Nrg1* (red), and DAPI (blue). Co-expressing cells are denoted by yellow arrowheads Scale bar= 50µm.

**Figure S6.**
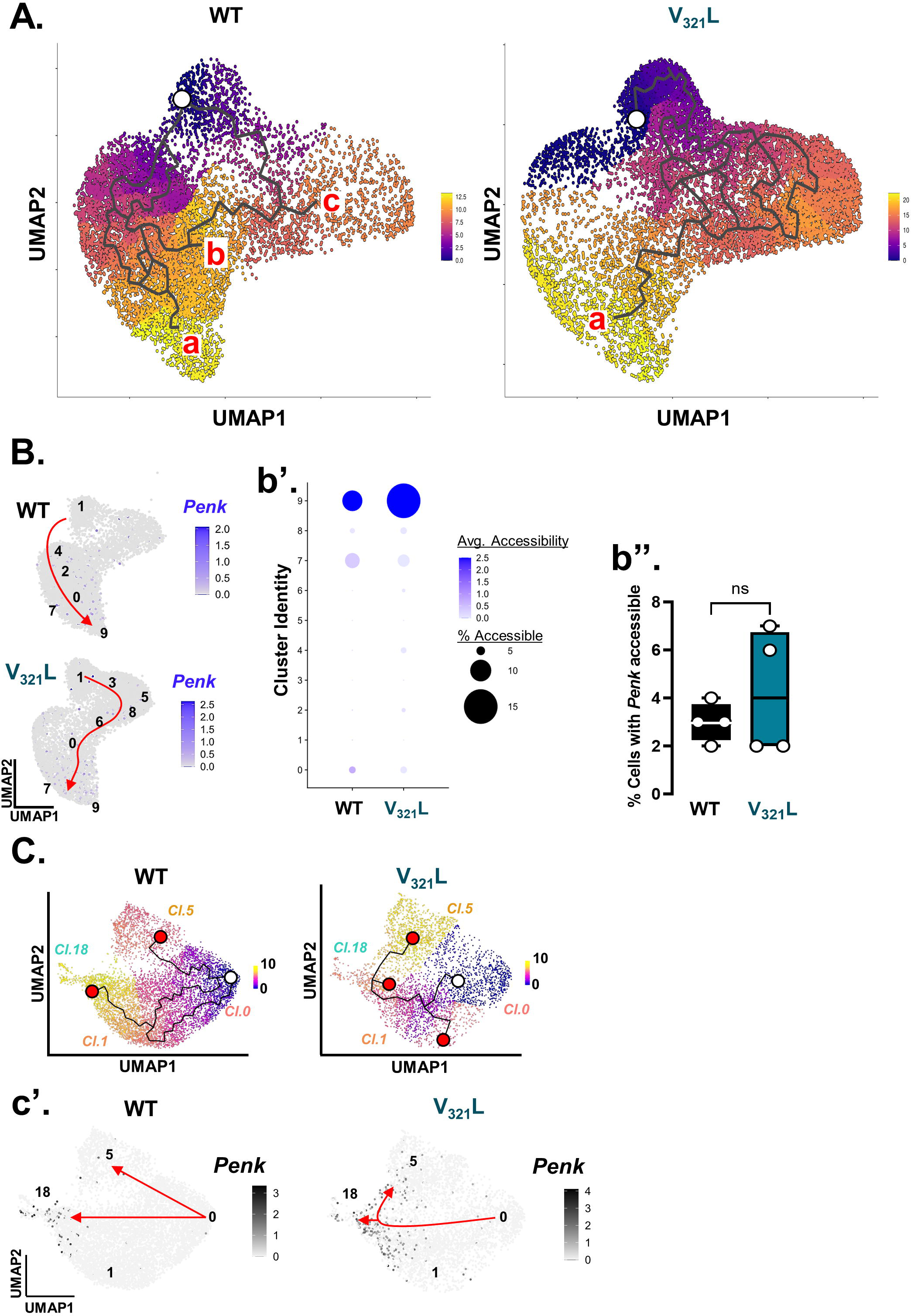
*Penk* gene accessibility increases along the GC maturation trajectory. **A.** UMAPs show snATACSeq data from WT (**Left**) and V_321_L (**Right**) mice with trajectories and pseudotime overlays indicated by the color according to the heatmap legend computed using Monocle3. End-trajectory points are labeled by “a” “b” and “c” in order of decreasing pseudotime. Note the dramatically different trajectories in WT vs. V_321_L cells ending in cluster 9 (SGC-enriched cluster). **B.** UMAPs show accessibility of the *Penk* gene locus in WT and V_321_L nuclei. Each dot represents a single nucleus color coded by the gene activity score (accessibility of the transcription start site and 1kb surrounding regions). Cluster IDs are overlaid on the UMAPs with a schematic representation of the end-to-end trajectories represented by red arrows. Note the distinct progression between WT and V_321_L cells; WT: cluster 1➔4➔2➔0/7➔9 whereas V_321_L: cluster 1➔3➔5/8➔6➔0➔9. **b’.** Dot plot showing average gene accessibility for *Penk* represented by the color of the dots and percentage of cells showing accessibility within a given cluster represented by the size of the dots. In WT DG, accessibility of the *Penk* locus increases along the maturation trajectory being undetectable in clusters 0-2 and gaining accessibility towards the end of the trajectory. In V_321_L DG, the locus is also accessible in clusters 6,4, and 2. **b’’.** Quantification of percentage of GCs with the *Penk* locus accessible in GCs from WT and V_321_L mice. There were no significant differences between the genotypes (Mann-Whitney Test p=0.9, U=7). N=4 mice/genotype. **C.** UMAPs showing snRNASeq data from WT (**Left**) and V_321_L (**Right**) mice with trajectories and pseudotime overlays indicated by the color according to the heatmap legend computed using Monocle3 (same as Figure 5B). **c’.** UMAPs show expression of the *Penk* gene in WT and V_321_L nuclei. Cluster IDs are overlaid on the UMAPs with a schematic representation of the end-to-end trajectories represented by red arrows highlighting a single main trajectory in the V_321_L DG similar to the ATAC-Seq data.

**Figure S7.**
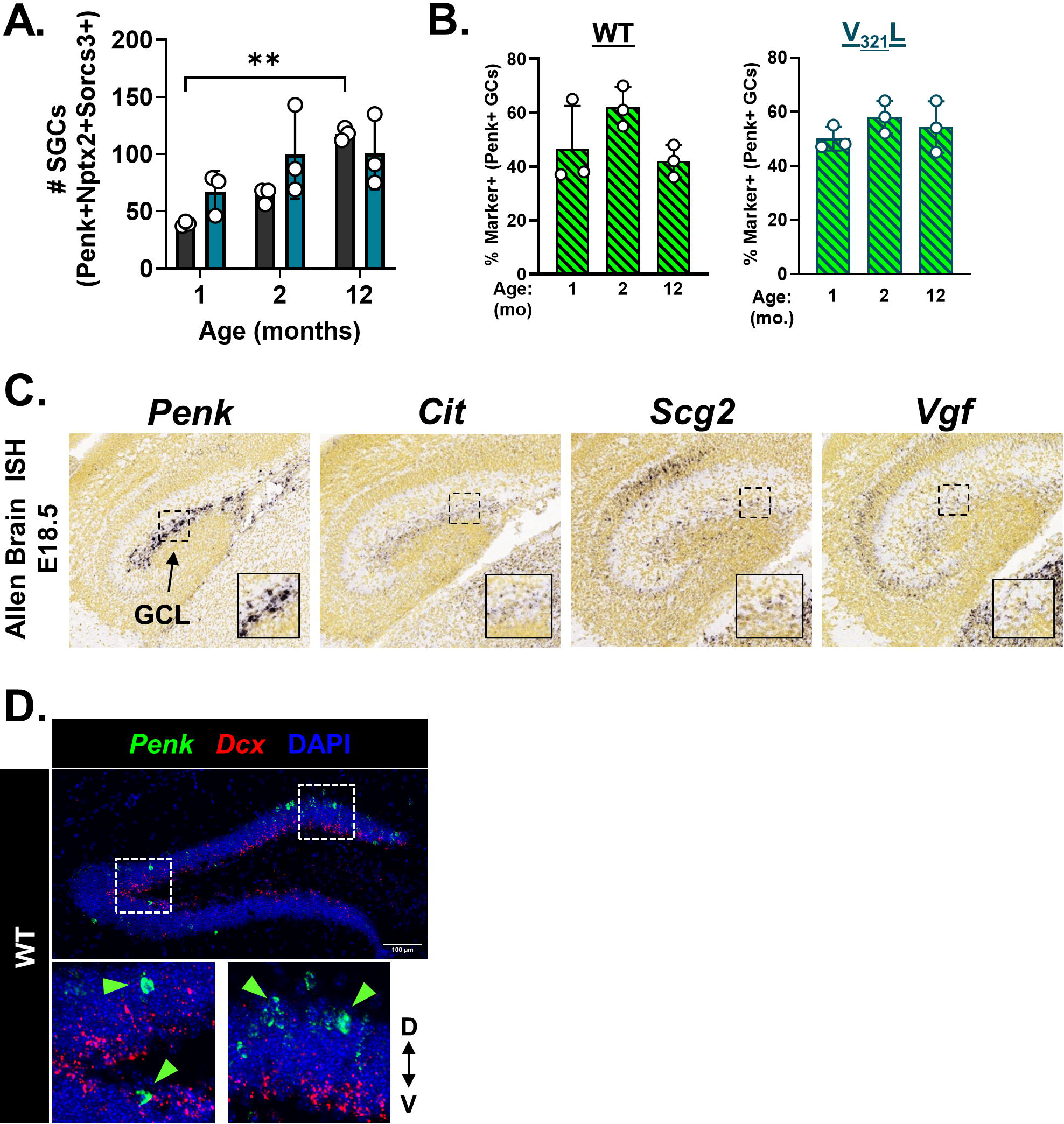
Age-dependent increase in SGC number is not accounted for by adult-born neurons. **A.** Age-dependent dynamics in SGC numbers are lost in the V_321_L DG. Quantification of total number of *Penk*+*Nptx2*+*Sorcs3*+ cells in the GCL of the suprapyramidal blade over age^$^ in the WT and V_321_L DG (Two-way ANOVA, effect of time p=0.008. Bonferroni corrected multiple comparisons-WT: 1mo. vs. 12mo. p=0.008, 2mo. vs. 12mo. p=0.06). ^$^Note that the 12 months old group comprised of animals aged 9-12 months of age. **B.** (**Left**) Quantification of *Penk+Nptx2+Sorcs3*+ (green bars with black diagonal stripes) cells as a percentage of all *Penk*+ cells over age^$^ (1-, 2-,and 12-months) in WT mice. There were no changes in the percentage of marker combination expressing cells over age (Brown-Forsythe ANOVA p=0.2, F= 2.1 (3, 4.75)). N= 3 mice/age group. (**Right**) Quantification of *Penk+Nptx2+Sorcs3*+ (green bars with black diagonal stripes) cells as a percentage of all *Penk*+ cells over age^$^ (1-, 2-,and 12-months) in V_321_L mice. There were no changes in the percentage of marker combination expressing cells over age (Brown-Forsythe ANOVA p=0.44, F= 0.99 (2.0, 4.3)). N= 3 mice/age group. ^$^Note that the 12 months old group comprised of animals aged 9-12 months of age. **C.** SGC-enriched genes are expressed in the embryonic DG with a spatial bias consistent with that of SGCs. Representative ISH images from the Allen Brain Atlas for *Penk*, *Cit*, *Scg2*, and *Vgf* in E18.5 DG. Insets show magnified images of the developing GCL and marker+ cells bordering the GCL. **D.** *Penk*+ cells in the GCL are not immature neurons. Representative image of a 2 month old WT DG showing ISH for *Penk* (green) and *Dcx* (red). Scale bar= 100µm. Insets show magnified images of two selected fields of view: *Penk* and *Dcx* transcripts were not found in the same cells despite some *Penk*+ cells occupying the bottom layers of the GCL populated by *Dcx*+ immature neurons. Green arrowheads indicate *Penk*+ GCs.

**Figure S8.**
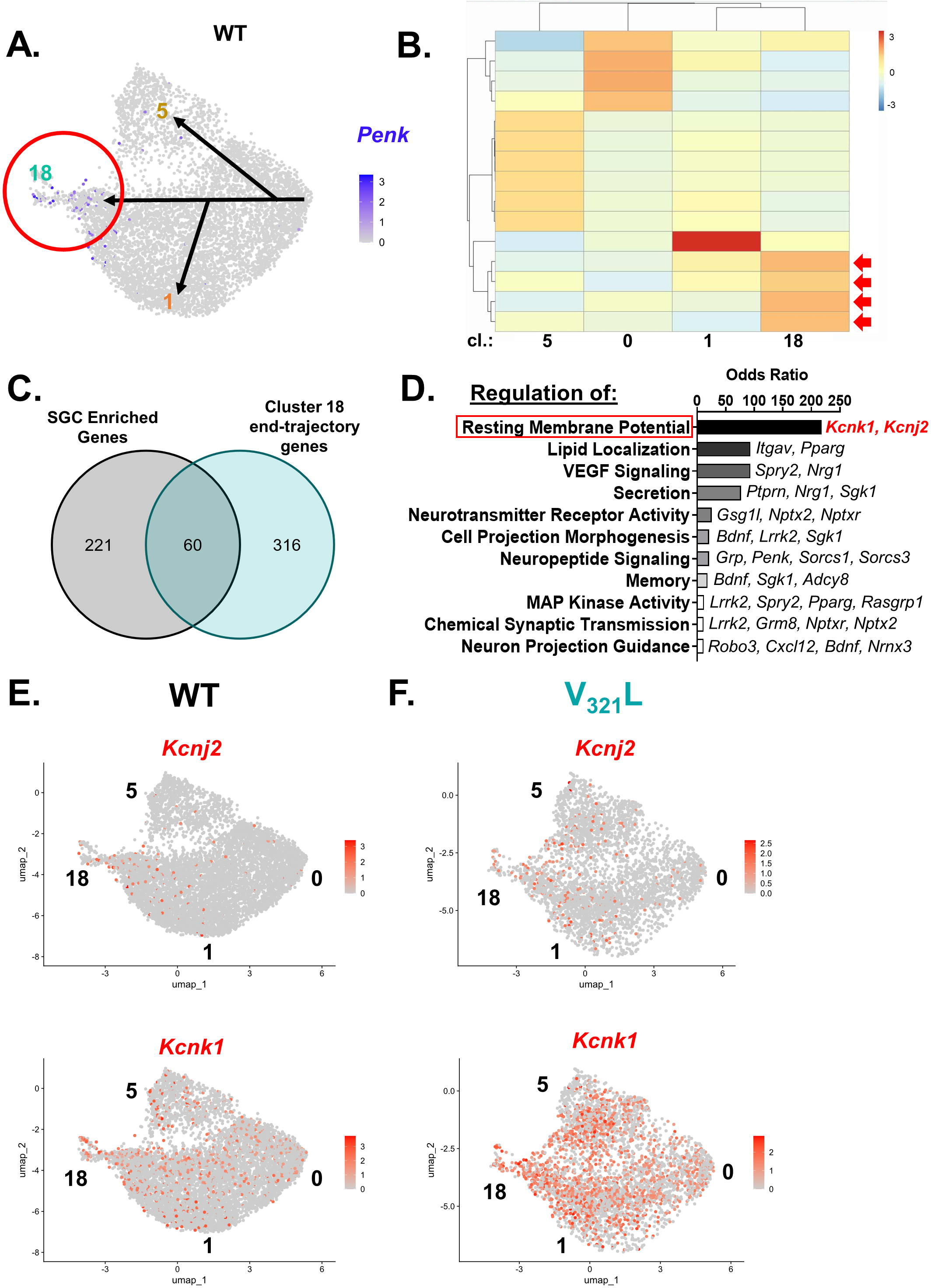
End-trajectory genes for the Penk+ GC-enriched cluster identify genes and phenotypes associated with SGCs. **A.** UMAP of *Penk* expression in the WT GCs indicated by the color according to the heatmap legend along with a schematic overlay of the different possible trajectories calculated by Monocle3. Trajectory ending in cluster 18, enriched for Penk+ GCs is highlighted by a red circle. **B.** Clustering of genes by pseudotemporal expression patterns resulted in several modules of genes associated with end-trajectories in different GC clusters. The heatmap shows individual gene modules as cells within the heatmap and the color represents the correlation of each module with end-trajectories in each of the clusters denoted below the heatmap. The color of the heatmap is according to the legend provided alongside. Red arrows indicate the four gene modules associated with end trajectories in cluster 18. **C.** An intersection set was created between genes enriched in expression in SGCs and genes part of the cluster 18 end-trajectory modules. 60 genes were found to be common between these two sets. These 60 genes were subjected to gene ontology analyses. **D.** Gene Ontology (GO) analysis for Biological Process. Different terms with statistical significance (adj.p<0.05) were selected and displayed by descending values of the odds ratio. Genes contributing to each term are displayed alongside the bar for each term. The top term-regulation of resting membrane potential is highlighted by a red box and the genes contributing to it are shown in red text. **E.** UMAPs of *Kcnj2* (**Top**) and *Kcnk1* (**Bottom**) expression in WT GCs indicated by the color according to the heatmap legend. Numbers alongside the UMAPs denote the cluster IDs. **F.** UMAPs of *Kcnj2* (**Top**) and *Kcnk1* (**Bottom**) expression in V_321_L GCs indicated by the color according to the heatmap legend. Numbers alongside the UMAPs denote the cluster IDs.

**Table S1.** Differentially expressed genes in Penk+ GCs vs. Penk-GCs.

**Table S2.** Genes correlated with pseudotime from Monocle3 trajectory analysis.

**Table S3.** ChEA & ENCODE enrichment for candidate transcriptional regulators of SGC-fate genes.

**Table S4.** DisGeNET enrichment for disease terms associated with SGC-fate genes.

**Table S5.** Differentially expressed genes in V_321_L vs. WT GCs.

**Table S6.** Differential motif activity in each ATAC cluster.

